# Genetic isolation by distance underlies color pattern divergence in red-eyed treefrogs (*Agalychnis callidryas*)

**DOI:** 10.1101/2021.05.21.445051

**Authors:** Meaghan I. Clark, Gideon S. Bradburd, Maria Akopyan, Andres Vega, Erica Bree Rosenblum, Jeanne M. Robertson

## Abstract

Investigating the spatial distribution of genetic and phenotypic variation can provide insights into the evolutionary processes that shape diversity in natural systems. We characterized patterns of genetic and phenotypic diversity to learn about drivers of color-pattern diversification in red-eyed treefrogs (*Agalychnis callidryas*) in Costa Rica. Along the Pacific coast, red-eyed treefrogs have conspicuous leg color patterning that transitions from orange in the north to purple in the south. We measured phenotypic variation of frogs, with increased sampling at sites where the orange-to-purple transition occurs. At the transition zone, we discovered the co-occurrence of multiple color-pattern morphs. To explore possible causes of this variation, we generated a SNP dataset to analyze population genetic structure, measure genetic diversity, and infer the processes that mediate genotype-phenotype dynamics. We investigated how patterns of genetic relatedness correspond with individual measures of color pattern along the coast, including testing for the role of hybridization in geographic regions where orange and purple phenotypic groups co-occur. We found no evidence that color-pattern polymorphism in the transition zone arose through recent hybridization. Instead, a strong pattern of genetic isolation by distance (IBD) indicates that color-pattern variation was either retained through other processes such as ancestral color polymorphisms or ancient secondary contact, or else it was generated by novel mutations. We found that phenotype changes along the Pacific coast more than would be expected based on genetic divergence and geographic distance alone. Combined, our results suggest the possibility of selective pressures acting on color pattern at a small geographic scale.

## Introduction

A central goal of evolutionary biology is to understand how phenotypic variation is generated, distributed, and maintained in natural systems (Mayr, 1963; Mitchell-Olds et al., 2007). Color pattern is a phenotype well-suited for evolutionary study because it is easily quantifiable and often has adaptive value (Cott, 1940; Orteu & Jiggins, 2020). Many compelling examples of natural selection in the wild have come from studying color-pattern variation across diverse empirical systems (e.g. Corl et al., 2010; Hoekstra et al., 2004; Lowry et al., 2012; Maan et al., 2008; Nachman et al., 2003; Pfeifer et al., 2018; Rosenblum et al., 2006; Streisfeld & Kohn, 2005; Supple et al., 2015; Twomey et al., 2015). In animals, color and color pattern can play important ecological roles, serving as, e.g., a deterrent to predation (crypsis, aposematism), as a signal to conspecifics (mate choice, territoriality), or both (Cummings & Crothers, 2013; Rojas, 2016; Selz et al., 2016; Stevens & Merilaita, 2009). Accordingly, color and color pattern are often shaped by selection, and variation in color traits can provide insight into the evolutionary dynamics that shape diversity. When present, color variation within a species can be distributed at a regional level with little variation within sites (polytypic), found at a single locality (polymorphic), or both (e.g., Wang & Summers, 2010). The distribution of color variation within species can provide particular insight on the spatial scale over which evolutionary dynamics act (Svensson, 2017).

A variety of evolutionary scenarios could explain phenotypic polytypism, polymorphism, and uniformity. For example, polytypic variation can arise and persist due to genetic drift, limited gene flow among populations due to environmental, geographic, or reproductive barriers, or selection against maladapted migrants (Endler, 1973; Lehtonen et al., 2009; Rosenblum et al., 2006, 2010). Polymorphism often occurs at transition zones between distinct morphs (Planes & Doherty, 1997) and can arise through novel mutations, contemporary gene flow, the retention of ancestral polymorphism, and/or ancient secondary contact (Lim et al., 2010; Roland et al., 2017). In such transition zones, hybridization could increase phenotypic variation within a population via the co-occurrence of distinct phenotypes unique to each parental population, or the generation of novel phenotypes in hybrids (Akopyan et al., 2020b; Anderson & Stebbins, 1954). Uniformity in color pattern is expected when there is substantial gene flow among morphs, or when color is under stabilizing selection (Duftner et al., 2006; Slatkin, 1985). Studying polytypism and polymorphism requires both georeferenced color-pattern data to quantify spatial patterns of phenotypic variation, and genotypes for those sampled individuals to understand the evolutionary processes generating those patterns of variation.

The striking color-pattern variation of red-eyed treefrogs, *Agalychnis callidryas* Cope 1862, provides an excellent system in which to study the mechanisms of extreme phenotypic variability. Red-eyed treefrogs exhibit color variation across their range from central Mexico to northern Colombia (Campbell, 1999; Savage & Heyer, 1967), with hypervariability in flank and leg color-pattern across populations in Costa Rica and Panama (Robertson & Robertson, 2008; Robertson & Zamudio, 2009). Flank and leg color are not sexually dimorphic, do not change within individuals in response to light intensity (Schliwa & Euteneuer, 1983), and are known to be heritable based on breeding studies that found intrapopulation crosses produce offspring with parental phenotypes (J.M. Robertson, unpublished data). Four distinct flank and leg color morphs exist in Costa Rica and Panama: blue, red/blue, orange, or purple (Robertson & Robertson, 2008). Across this part of the range, red-eyed treefrogs are polytypic, but generally not polymorphic (Robertson & Robertson, 2008; Robertson & Vega, 2011), except in a contact zone on the Caribbean slope of Costa Rica (Akopyan et al., 2020b). Flank and leg color-pattern likely evolve through both sexual selection due to preference for local morphs during mate choice (Akopyan et al., 2018; Jacobs et al., 2016; Kaiser et al., 2018) and natural selection for color pattern to potentially act as a warning signal of unpalatability to predators (Clark, 2019; Davis et al., 2016; Robertson & Greene, 2017). Thus, the evolutionary dynamics and impacts of selection have likely shaped the polytypic and polymorphic distributions of color pattern in red-eyed treefrogs.

We focused on populations of red-eyed treefrogs from mainland and peninsular populations of the Pacific slope of Costa Rica, where color variation is characterized by a north-south gradient. In the north, frogs generally have orange flanks and legs (orange morph), while in the south, they display purple flanks and legs (purple morph, Fig. 1). Recent field sampling efforts identified polymorphic sites where color pattern transitions from the orange to the purple morph (Central 2, Fig. 1). Previous research revealed a complex relationship between geography, color pattern, and mitochondrial and nuclear markers (Robertson et al., 2009). Within the Pacific slope, northern and southern sites do not share mitochondrial haplotypes, except at a single centrally located site (Robertson and Zamudio 2009; Fig. 1, Site 4; Table 1). We aim to understand the evolutionary processes underlying both polytypic and polymorphic color-pattern distributions on the Pacific coast.

**Figure 1.**
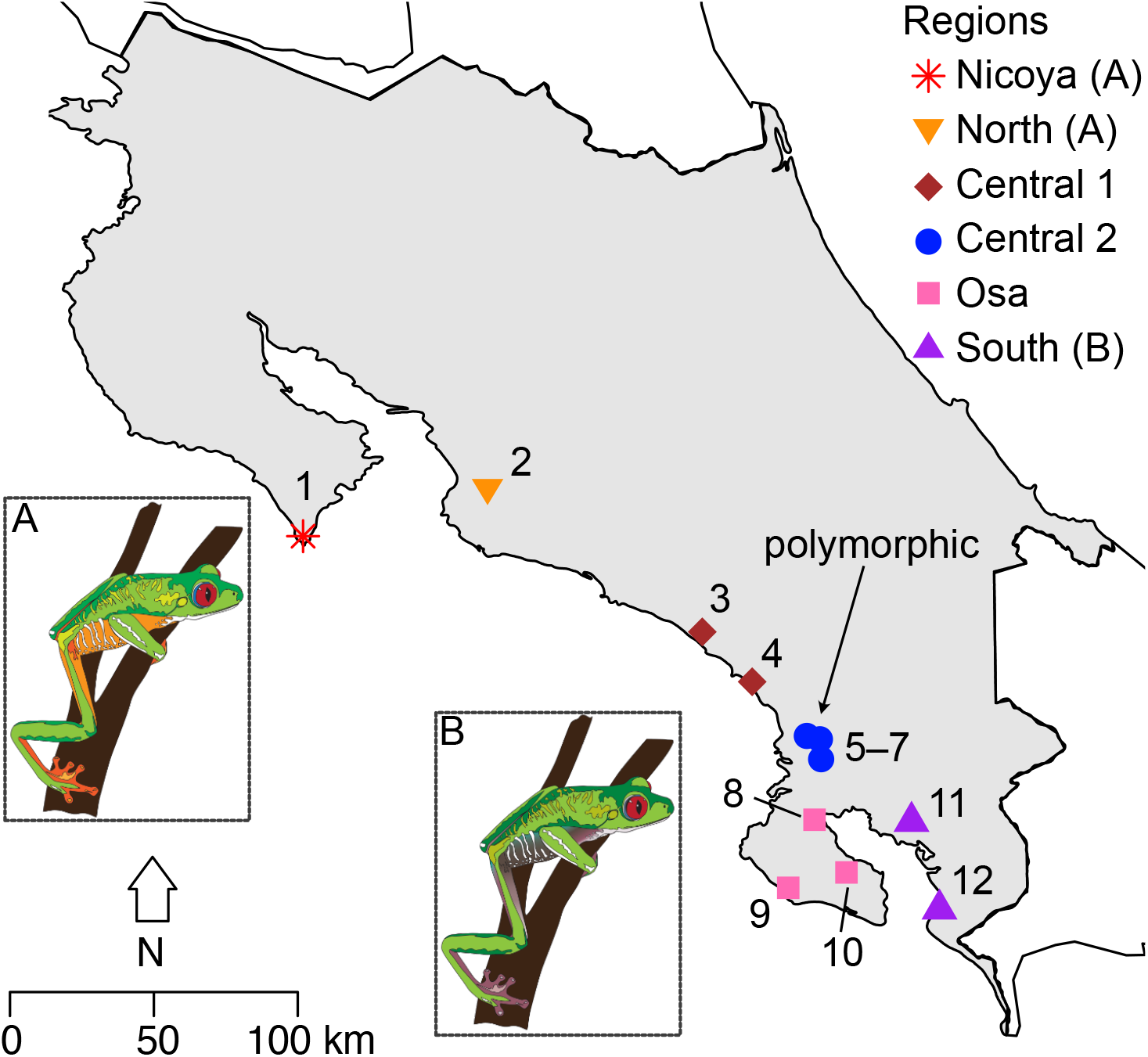
Map indicating the 12 sampling sites for red-eyed treefrogs (*Agalychnis callidryas*) along the Pacific coast of Costa Rica. Each site is assigned to one of six biogeographic regions: Nicoya, north, central 1, central 2, Osa, and south. Arrow indicates polymorphic sites. Insets A and B: red-eyed treefrog (*Agalychnis callidryas*) color morphs, drawn from photographs, illustrate color patterns typical of northern orange (A, Nicoya, North) and southern purple (B, South) regions (illustrations: Cynthia J. Hitchcock).

**Table 1.** Location, geographic coordinates, sample size, mtDNA clade (Robertson & Zamudio, 2009) and genetic diversity for 12 sampling sites of red-eyed treefrogs (*Agalychnis callidryas*) along the Pacific coast of Costa Rica. MtDNA haplotype not identified (––) for some sites. Estimates of nucleotide diversity (π) and Wu and Watterson’s theta (θ_w)_ based on RADseq dataset. See Fig. 1 for region designations.

| # | Site | Province | Region | Latitude (N) | Longitude (W) | *n* | mtDNA clade | $\pi_p$ | $\theta_w$ | Private alleles |
| --- | --- | --- | --- | --- | --- | --- | --- | --- | --- | --- |
| 1 | Cabo Blanco | Guanacaste | Nicoya | 9.5805 | -85.1246 | 9 | B | 0.088 | 0.071 | 907 |
| 2 | Bijagual | San Jose | North | 9.7256 | -84.5313 | 9 | B | 0.154 | 0.131 | 213 |
| 3 | Firestone | Puntarenas | Central 1 | 9.274927 | -83.8589 | 3 | — | 0.167 | 0.126 | 0 |
| 4 | Uvita | Puntarenas | Central 1 | 9.1235 | -83.7011 | 5 | A, B | 0.165 | 0.149 | 3 |
| 5 | Cortes | Puntarenas | Central 2 | 8.95517 | -83.52177 | 12 | — | 0.169 | 0.152 | 66 |
| 6 | Palmar Sur | Puntarenas | Central 2 | 8.94488 | -83.48444 | 20 | — | 0.168 | 0.144 | 104 |
| 7 | Sierpe | Puntarenas | Central 2 | 8.8892 | -83.477 | 17 | A | 0.174 | 0.163 | 34 |
| 8 | El Campo | Puntarenas | Osa | 8.6909 | -83.5013 | 17 | A | 0.163 | 0.144 | 83 |
| 9 | Sirena | Puntarenas | Osa | 8.480261 | -83.59012 | 19 | — | 0.165 | 0.145 | 80 |
| 10 | El Tigre | Puntarenas | Osa | 8.52678 | -83.40053 | 19 | — | 0.165 | 0.143 | 52 |
| 11 | Gamba | Puntarenas | South | 8.69333 | -83.19522 | 19 | — | 0.164 | 0.143 | 172 |
| 12 | Pavones | Puntarenas | South | 8.4204 | -83.1069 | 25 | A | 0.161 | 0.138 | 173 |

In this study, we document the distribution of polytypic and polymorphic populations across the orange-purple color-pattern gradient and combine those data with a genomic dataset generated using RAD-seq to understand the evolutionary mechanisms that generate and maintain patterns of diversity. We expand sampling from previous studies to examine genomic ancestry and phenotypic patterns across and within orange, purple, and polymorphic sites. We first address two hypotheses regarding polytypic patterns across the entire study area: (1A) purple and orange morphs represent distinct genetic demes, consistent with selection for reproductive isolation among these morphs (Akopyan et al., 2018), or alternatively, (1B) samples along the coast are characterized by a pattern of genetic isolation by distance (IBD) that is not correlated with phenotypic variation, which could indicate local selection acting to maintain color morphs (here we refer to IBD as a pattern of increasing genetic divergence with geographic distance). We investigate the relationships between phenotype and geographic distance, latitude, and genetic diversity using Mantel tests and Bayesian models.

Focusing on the transition zone, we then combine analyses of color pattern, genetic variation, and relatedness to investigate the phenotypic and genomic profile of polymorphic sites. First, we characterize the distribution of color phenotypes at polymorphic sites using principal components analysis and discriminate function analysis to ask if (2A) only orange and purple morphs are present, or (2B) unique color phenotypes are present alongside orange and purple morphs. We then use genomic clustering analyses to examine the genomic makeup of phenotypically polymorphic populations compared to other (non-polymorphic) populations: (3A) If there is little or no gene flow between orange and purple morphs, clustering analyses will show distinct parental groups at polymorphic sites. (3B) Alternatively, if there is gene flow between morphs, clustering analyses will show individuals at polymorphic sites assigned to the same genetic cluster, despite distinct phenotypes. Next, we use the program entropy (Gompert et al., 2014) to determine if (4A) hybridization underlies polymorphism, or if (4B) polymorphism has been generated through another mechanism (i.e., retained from an ancestral population or ancient secondary contact, and/or novel morphs have been generated by new mutations). By integrating analyses of variation at both large (transect from north to south, including polytypic sites and transition zone) and small spatial scales (polymorphic sites in transition zone), we can test hypotheses about how color pattern evolves at microgeographic scales.

## Methods

### Field sampling and study sites

Our study focused on 12 sites along the Pacific coast of Costa Rica, west of the continental divide (Fig. 1). Pacific red-eyed treefrogs are isolated from Caribbean populations by the Cordillera de Talamanca (Robertson & Zamudio, 2009). The Pacific slope is geographically and environmentally complex: two peninsulas, the Nicoya and Osa, are potentially isolated from mainland sites through geographic barriers and inhospitable habitat (Robertson et al., 2009). Previous studies characterized red-eyed treefrogs at northern sites (Sites 1–8) as orange and at a southern site as purple (Site 12) (Robertson & Vega, 2011). Here, we expanded previous geographic sampling (Robertson et al., 2009) by adding six sites to assess fine-scale variation in phenotype and genotype, including: the central coast (Site 3), the Osa Peninsula (Sites 9 and 10), mainland adjacent to Osa Peninsula (Sites 5 and 6), and Gulfo Dulce (Site 11). We included 9– 25 individuals from previously sampled sites, and 3–20 individuals from new sites (Table 1). We bred frogs from Site 2 with frogs from Site 12 in captivity at California State University, Northridge (IACUC 1819-005) to produce F1 hybrids to serve as controls in ancestry analyses (see genomic analyses below). Unfortunately, these frogs did not survive until adulthood, so we were unable to assess their phenotype.

We grouped sites together into six regions based on potential geographic barriers, and divisions between known phenotypic and genetic groups (Fig. 1). The Nicoya Peninsula (Site 1) is geographically isolated from all mainland populations by dry forest, which is an unsuitable habitat for red-eyed treefrogs. The Central 1 (Sites 3-4) region is separated from the North (Site 2) based on distinct mtDNA haplotypes (Robertson & Zamudio, 2009). The Central 2 region (Sites 5-7) contains both orange and purple frogs whereas frogs in the South (Sites 11-12) only have purple legs. The Terraba and Sierpe Rivers separate the Osa Peninsula (Sites 8–10) from the mainland (Kohlmann et al., 2002; Robertson & Vega, 2011).

At each site, we captured frogs by hand and took digital photographs with a black-white-gray standard (QPcard 101 or 102, Adorama Camera Inc., New York, New York) of the posterior aspect of their legs (Suppl. Table 1, for samples collected before 2015, see Robertson & Robertson 2008, Robertson & Vega 2011; for samples collected in 2015 see Akopyan et al. 2020b). We obtained a toe clip for genomic analyses, which was stored in 100% ethanol. Frogs were released within one day at the site of capture. We also took toe clips from eight lab-bred F1 hybrids.

### Quantifying color pattern

Photos were color corrected in Photoshop CC v. 18.0.0 (Adobe, San Jose, CA) using the QPcard 101 and QP102 gray standard (same subset of standards in both cards). To quantify color pattern, we focused on the posterior leg because leg and flank color-pattern are strongly correlated (Robertson & Robertson, 2008). The colored region of the frogs’ legs can be either uniform in color or contain a gray/blue patch on the inner leg (Suppl. Fig. 1). In all cases, we first excluded the patch and selected the colored region of one leg and used the “average” function in Photoshop to measure hue, saturation, and brightness (Adobe, San Jose, CA). Because orange and purple have similar hue values, the saturation and brightness values were useful to distinguish color pattern within and among populations (Robertson & Robertson, 2008). We then used ImageJ (Abràmoff et al., 2005) to quantify the percent of pixels on the leg that were covered by the gray patch, which was easily distinguished from the dominant portion of the leg using saturation values.

We used principal component analysis (PCA) in R v. 3.5.3 (R Core Team 2016) based on hue, saturation, brightness, and percent gray of the hind legs to visualize the distribution of color-pattern variation among red-eyed treefrog populations within and among regions. Because it is likely that different loci underly color and pattern in red-eyed treefrogs, we also used PCA to visualize the distribution of color measurements separate from pattern. Photographs were taken with different cameras (Canon PowerShot SX160 and Nikon Coolpix 5700), but we do not see evidence of batch effects corresponding to camera type (Suppl. Fig. 2, 3). We conducted a linear discriminant function analysis (DFA) using “lda” function in the MASS v. 7.3-51.4 package in R v. 3.5.3. to test whether individuals were correctly assigned to their site of origin based on either hue, saturation, brightness, and percent gray. Individuals were classified to the site with the highest posterior probability. We conducted this analysis with all measurements and each measurement separately.

### Molecular methods

DNA was extracted from frog toe clips using a DNeasy Blood and Tissue kit (Qiagen, Valencia, CA) per the manufacturer’s protocol, except as described below. We eluted DNA in 50 uL AE to increase concentration. After extraction, 4 uL of 1 mg/mL RNAse A was added to each sample to remove RNA contamination. DNA concentration was measured using a Qubit 2.0 Fluorometer (ThermoFisher, Waltham, MA). Restriction site associated DNA sequencing (RAD-seq) libraries (Ali et al., 2016; Etter et al., 2011) containing samples from 12 sites and eight lab-bred F1 hybrids were prepared with restriction enzyme sbf1. We included 4 to 20 individuals per site (Table 1). Libraries were sequenced on one lane of a HiSeq4000 (PE 150; Illumina San Diego, California).

### Quality filtering and SNP calling

We processed RAD-seq data as in Akopyan et al. (2020b). Briefly, raw reads with the cut site on the reverse read were rescued using a custom Perl script. The *clone_filer* program in STACKS v. 1.34 (Catchen et al., 2013) was used to remove PCR clones. Sequence data were demultiplexed using the *process_radtags* function in STACKS. Reads containing base pairs with a phred score less than 10 were removed. We used *ustacks* in STACKS for *de novo* assembly, requiring a minimum of three reads to create a stack, and a maximum of four base-pair differences between stacks. A pseudo-reference genome was created using a custom Perl script. Over half of the individuals had to have a stack for it to be used in the pseudo-reference genome. Reads were aligned to the pseudo-reference genome using “aln” and “samse” in the program BWA v. 0.7.15-r1142 (Li & Durbin, 2009). We set the maximum edit distance to four, seed to 20, maximum edit distance within the seed to two, and the read trimming parameter to 10. Samtools v. 1.9 (Li et al., 2009) was used to index, sort, and merge alignments. We used the Bcftools v. 1.3.1 “call” function to call variants and create a VCF file (Li, 2011). We filtered the data to remove low coverage loci (<2x per individual) and to retain a single SNP per contig. Loci with mean allele frequencies <0.5% and individuals with >70% missing data were not included in subsequent analyses. Pipeline and scripts are available online (Akopyan et al., 2020a).

### Population genomic analyses

We calculated two measures of relative genetic variability, pi at polymorphic sites (π_*p*_), and Wu and Watterson’s theta (θ_w_, Watterson 1975) at polymorphic sites, in R using custom scripts. We defined *S* as the total number of segregating sites in the global dataset, and *s*_*i*_ as the number of those sites genotyped in the *i*th individual. To calculate π_*p*_, we first calculated heterozygosity at polymorphic sites for each individual; in the *i*th individual, this is the number of sites at which that individual was called heterozygous divided by *s*_*i*_. To calculate π_*p*_ for a sampling site, we then averaged the heterozygosities of all individuals within that site (Charlesworth & Charlesworth, 2010). Wu and Watterson’s theta for a population was calculated as the number of polymorphic sites in that population divided by the product of *S* and *a*_*k*_, where *a*_*k*_ is Watterson’s correction factor for the number of individuals in that population (Charlesworth & Charlesworth, 2010; Watterson, 1975). We calculated the number of private alleles in each population in R. We used a Bayesian clustering method implemented in the program entropy (Gompert et al., 2014) to estimate admixture proportions and interpopulation ancestry. We excluded sites 1 and 8–10 from the entropy analysis to limit the dataset to putative parental and hybrid groups based on admixture proportions. To assess the possibility of hybridization across sites, we plotted interpopulation ancestry (Q_12_) against admixture proportion (q) (Gompert & Buerkle, 2016).

Interpopulation ancestry, the proportion of an individual’s genome where each allele at a locus is from a different putative parental group, separates out F1 hybrids (Q_12_ near 1), multigenerational hybrids (intermediate Q_12_) and individuals with late-stage hybrid genomes (Q_12_ near 0). The lab-reared F1 hybrids served as a useful control for what we expect for F1 hybrids. Multigenerational hybrids were expected to show elevated interpopulation ancestry, but below that of F1 hybrids.

We ran Mantel tests in R package ade4 v.1.7-13 (10,000 replicates) to test for correlations between geographic distance and genetic distance; geographic distance and phenotypic distance; and genetic and phenotypic distance (Chessel et al., 2004). We assumed frogs would avoid crossing bodies of saltwater, so we calculated geographic distances among populations as over-land distances. We calculated pairwise *F*_ST_ between sites in the R package BEDASSLE v. 1.5 (Bradburd et al., 2013; Weir & Hill, 2002). We estimated phenotypic distance between sites by calculating the pairwise Euclidean distance of average hue, saturation, brightness, and percent gray for each site. We also tested for relationships between each phenotypic measurement and geographic/genetic distance individually. A *Q*_ST_–*F*_ST_ or *P*_ST_–*F*_ST_ approach is not justified for this study because we did not measure phenotype in a common garden, and the genetic basis of color pattern in red-eyed treefrogs is unknown (Pujol et al., 2008).

Because we observed a strong pattern of genetic IBD (see Results), we used the R package conStruct v.1.0.4 to assess evidence for discrete genetic clusters while accounting for continuous differentiation resulting from geographic distance (Bradburd et al., 2018). The spatial model in conStruct estimates admixture proportions for each sample while simultaneously accounting for the decay in genetic relatedness with distance within each discrete group. We ran four chains for each spatial and non-spatial model specifying a number of discrete groups (*K*) between 1 and 6. We selected the chain with the highest mean posterior probability and assessed the biological importance of each added *K*-value by (1) comparing the contribution of each layer to overall covariance, (2) cross-validation analysis that compares the predictive accuracy of models using eight replicates and 1000 iterations, and (3) assessing the biological realism of the model given previous knowledge of the system. To explore genetic clustering in restricted spatial scales, we repeated conStruct analyses for three subsets of the data: without Site 1 (“Without Nicoya”), northern sites (Sites 1 and 2), and southern sites (Sites 5-12).

To evaluate phenotypic patterns while accounting for genetic relatedness, we fit three sets of Bayesian linear models implemented in rstan v.2.12.2 (Stan Development Team, 2018): (1) we modeled individual phenotype measurements (hue, saturation, brightness, and color-pattern PC1) each as a function of latitude, (2) we modeled the average phenotype for each site as a function of latitude, and (3) we modeled phenotypic variation at a site as a function of latitude, and, separately, of θ_w_. All models included a covariance structure that was a function of the sample covariance of allele frequencies between individuals and were run four times, for 2000 iterations each. These models allowed us to detect variation in phenotype above and beyond that explained by covariance at putatively neutral genetic markers.

## Results

### Phenotypic variation

Phenotypic patterns along the Pacific coast matched previous studies: frogs had orange legs and flanks in the northern regions and purple flanks and legs in the south. We confirmed that the three sites in the Central 2 region were polymorphic. The first two principal components in our color-pattern PCA explained 75.6% of the total variance in measured leg color-pattern metrics (Fig. 2, Suppl. Fig. 4). Sites in the Nicoya, North and Central 1 regions (orange legs) formed one cluster in phenotypic space, and sites in the South (purple legs) formed another. Sites in the Central 2 region were generally intermediate, but closer to the orange cluster. Osa sites were variable: Site 10 clustered with the purple/south sites, Site 8 clustered with the orange/northern sites, and Site 9 was intermediate. Color pattern PCs 2 and 3 are visualized in the Supplemental Materials (Suppl. Fig. 5). The first two principal components in the color-only PCA explained 84.3% of the total variance in the leg coloration metrics. The relative location of sites in the color-only PCA was similar to the color-pattern PCA, but there was less overlap between sites (Suppl. Fig. 6). The percent of the hind leg taken up by the inner gray patch increased from 0% in the Nicoya and North regions to an average of 33.3% in the South region (Suppl. Fig. 7).

**Figure 2.**
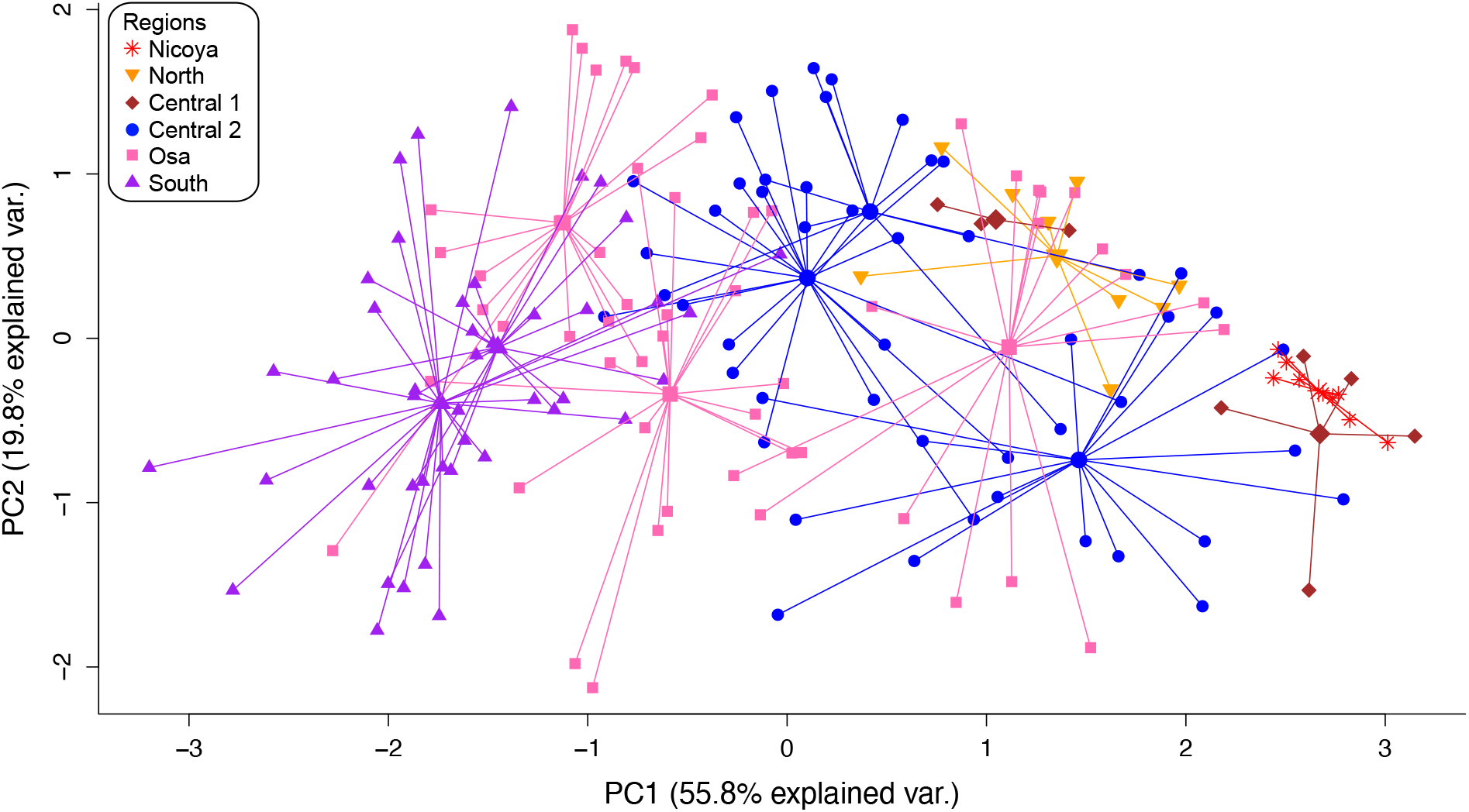
Principal component analysis of leg phenotype for red-eyed treefrogs (*Agalychnis callidryas*) sampled from 12 sites along the Pacific coast of Costa Rica. Lines connect individuals (small symbol) with the mean phenotype of each sampling site (large symbol). Individuals and means are color coded by region.

The DFA run with all phenotypic measurements had an overall predictive accuracy of 69.0%. Individuals from the Nicoya had 100% correct assignment (Table 2). The majority of frogs were correctly assigned to their region of origin, with the exception of orange frogs, which were frequently assigned to phenotypically similar sites in different regions (Table 2). Frogs from polymorphic sites in the Central 2 region had high misclassification probabilities (Table 2). Most (72.7%) purple individuals from the South were assigned to the correct region. Over half of the individuals from the Osa were assigned to the correct site; misclassified individuals were most often assigned to sites with color patterns similar to their respective Osa site (e.g., 26% of purple morph frogs from Site 10 were misassigned to Site 11, another purple morph site). Overall predictive accuracy decreased in DFA performed using single phenotypic measurements (hue: 31.0%, saturation: 30.5%, brightness: 23.0%, gray patch percentage: 19.5%, Suppl. Tables 2-5).

**Table 2.**
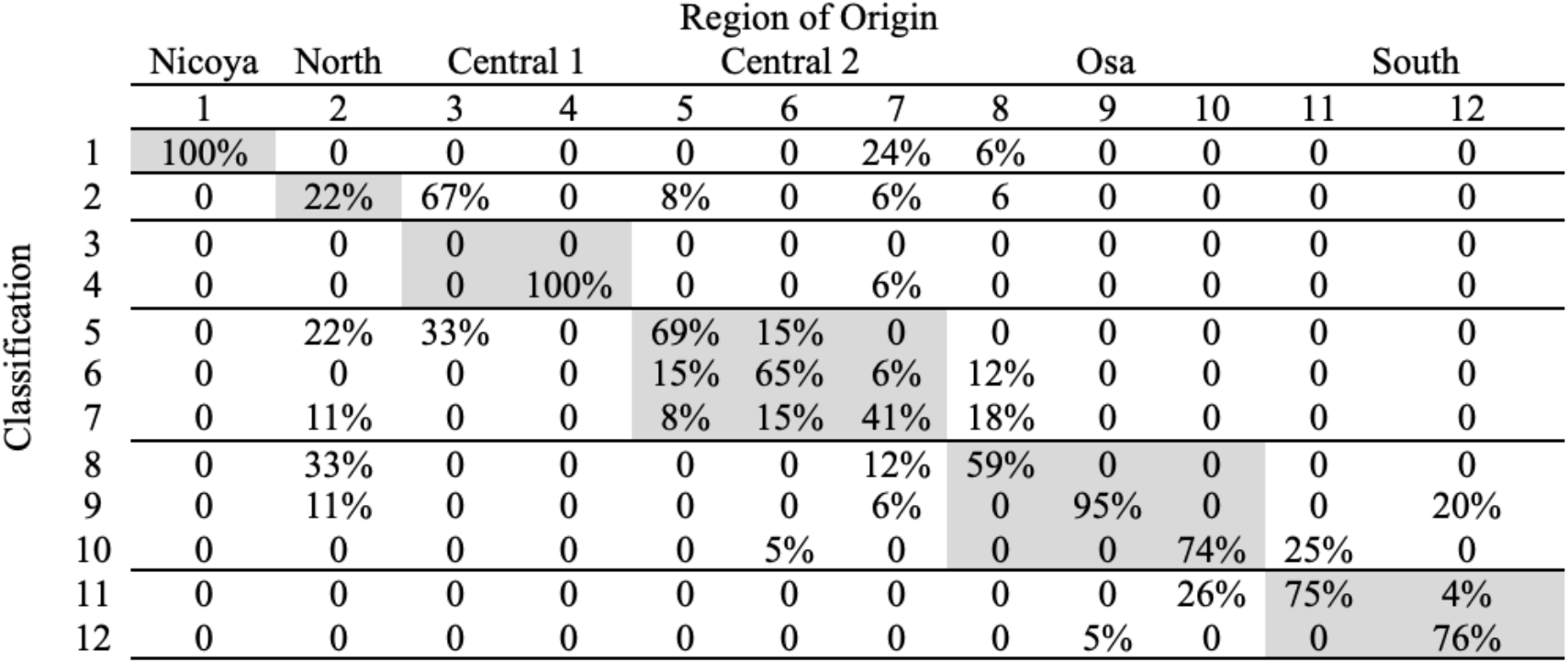
Discriminant function analysis of leg color pattern for 12 sampling sites of red-eyed treefrogs (*Agalychnis callidryas*) along the Pacific coast of Costa Rica. Each individual was classified based on its leg color pattern and assigned to a sampling site.

### Genetic variation

After filtering, we retained 71,746 SNPs in 174 individuals. The mean coverage per SNP was 9.3X with a standard deviation of 2.97. Descriptive statistics indicated similarity in genetic diversity among sites, except for Site 1, which had lower genetic diversity (Table 1). Site 1 also had the highest number of private alleles (907, Table 1). The highest genetic diversity was found at Site 7 in the Central 2 region. Pairwise *F*_ST_ values ranged from 0.025 to 0.422, with the highest value between Sites 1 and 3 (Table 3). The genetic PCA revealed genetic clustering corresponding to geographic regions (Fig. 3, Suppl. Fig. 8). Combined, the first two principal components explained 12.6% of variation. The Nicoya, North, Central 1, the Osa, and the lab-generated hybrids were genetically distinct, while South and Central 2 regions clustered together. Genetic PCs 2 and 3 are visualized in the Supplemental Materials (Suppl. Fig. 9).

**Table 3.**
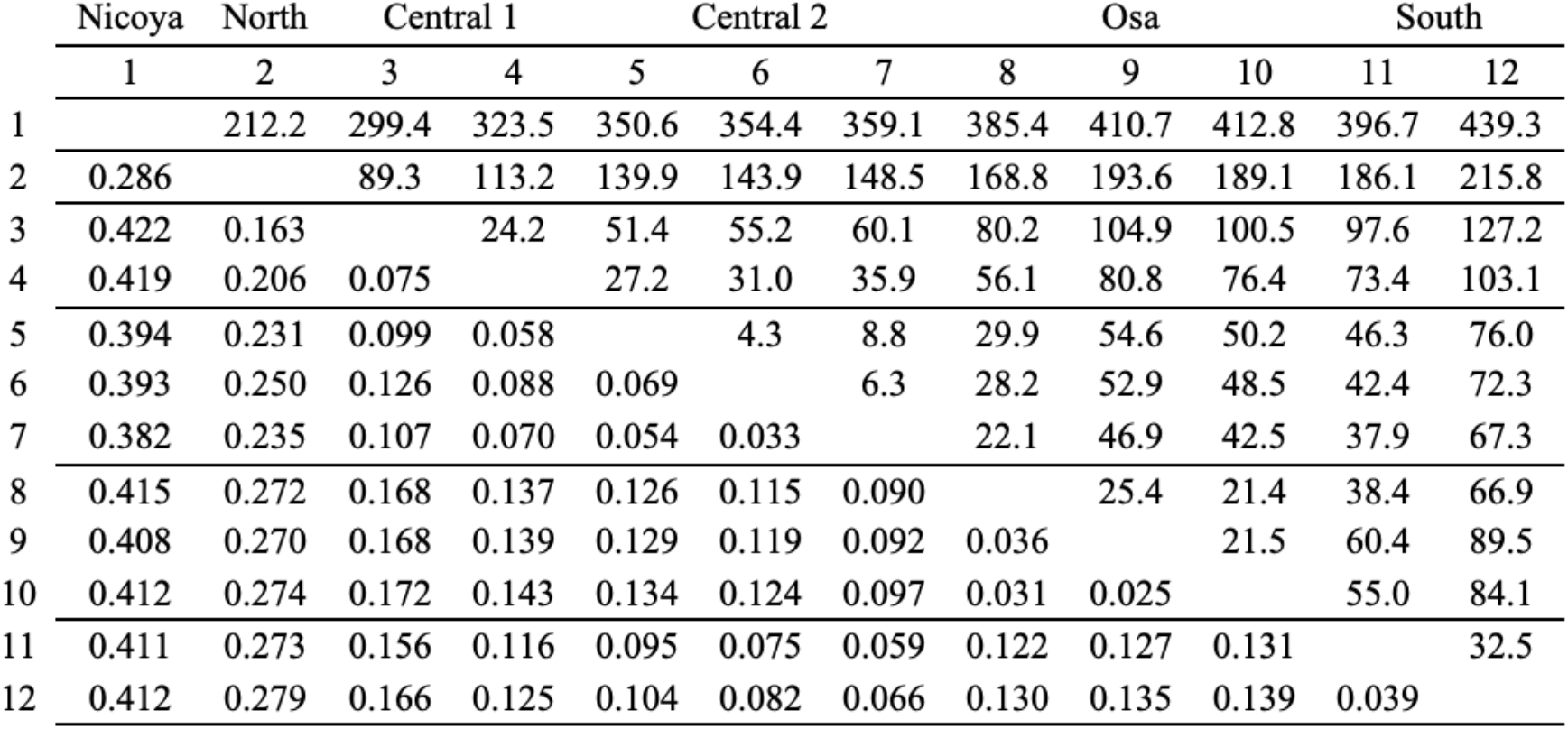
Pairwise *F*_ST_ (below diagonal) and geographic distance in kilometers (above diagonal) for 12 sampling sites along the Pacific coast of Costa Rica. Lines denote regional boundaries.

**Figure 3.**
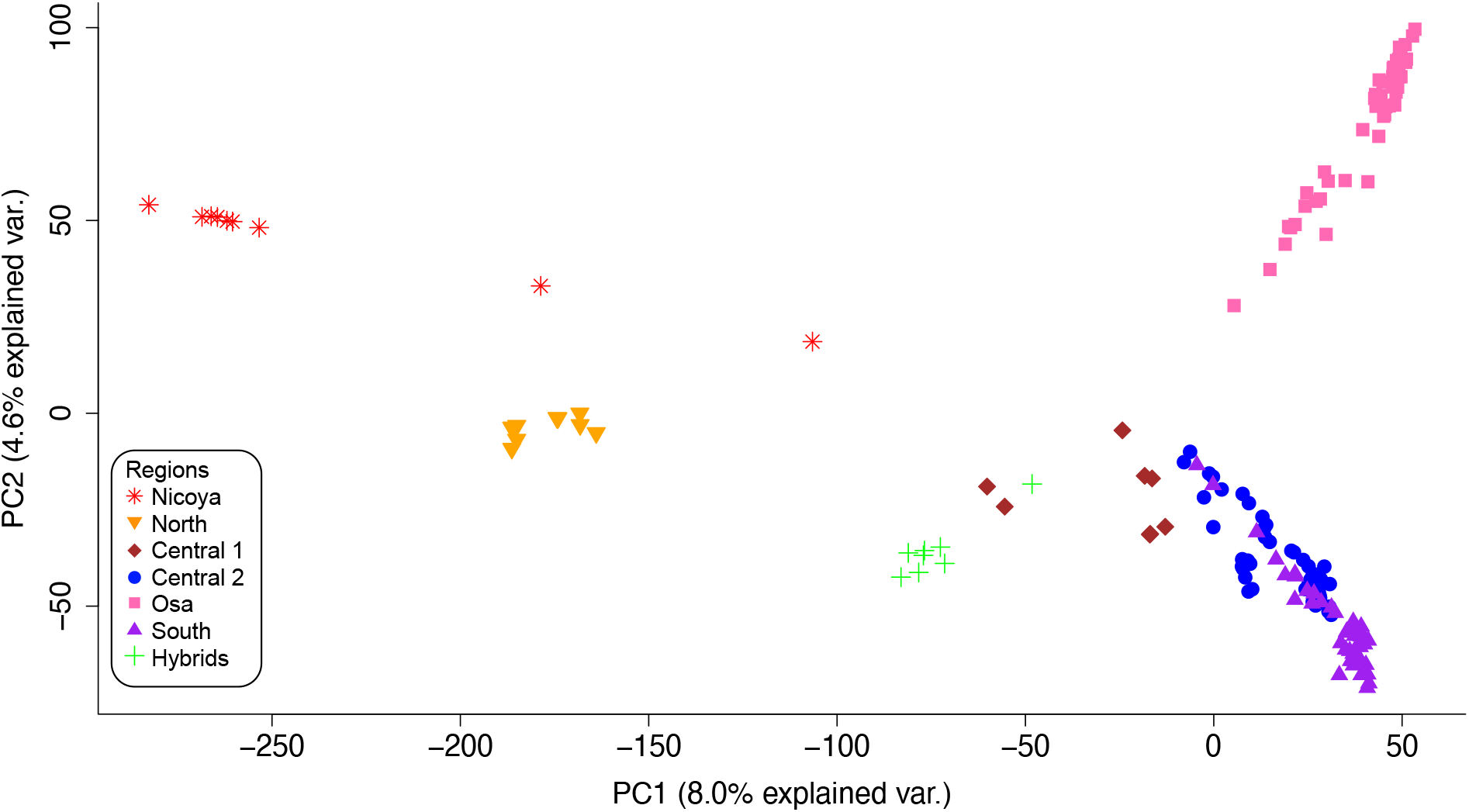
Principal component analysis of genomic SNPs for red-eyed treefrogs (*Agalychnis callidryas*) sampled from 12 sites along the Pacific coast of Costa Rica. Symbols represent individuals. Both type of symbol and color represent region. Green plus symbols are lab generated hybrids between Site 2 and Site 12.

The comparison of admixture proportion (q) and interpopulation ancestry (Q_12_) from entropy showed no evidence of recent hybridization when parental populations were assigned as North and South. Lab-generated hybrids had intermediate admixture proportions and high interpopulation ancestry scores, as expected. No wild frogs had elevated interpopulation ancestry scores (Fig. 4).

**Figure 4.**
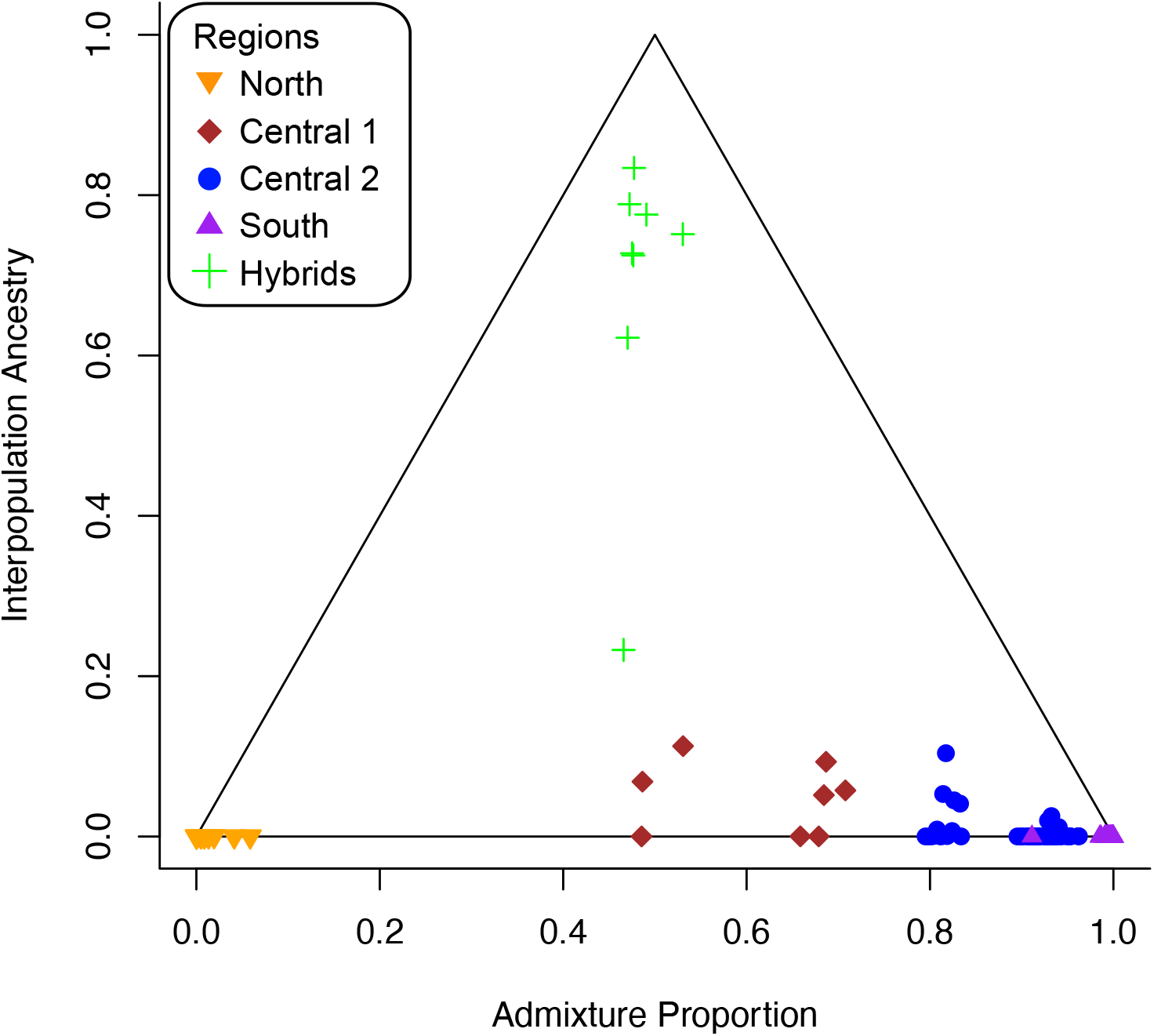
Scatterplot of admixture proportion (q) and interpopulation ancestry (Q_12_), the proportion of an individual’s genome where one allele is assigned to each parental group, for red-eyed treefrogs (*Agalychnis callidryas*) sampled from 5 sites along the Pacific coast of Costa Rica. Symbols represent individuals and are color coded by region. Only lab generated hybrids (green) show high interpopulation ancestry indicative of hybrid origin.

### Genotypic and phenotypic patterns

Mantel tests revealed a strong positive relationship between geographic and genetic distance (*r* = 0.965, *p* <0.001) and weak positive relationships between geographic and phenotypic distance (Mantel test, *r* = 0.336, *p* = 0.0085) and genetic and phenotypic distance (Mantel test, *r* = 0.219, *p* = 0.106, Fig. 5). Results between composite phenotypic distance and individual phenotypic measurements were consistent, and results for individual phenotypic measurement tests are presented in the Supplemental Materials (Suppl. Fig. 10).

**Figure 5.**
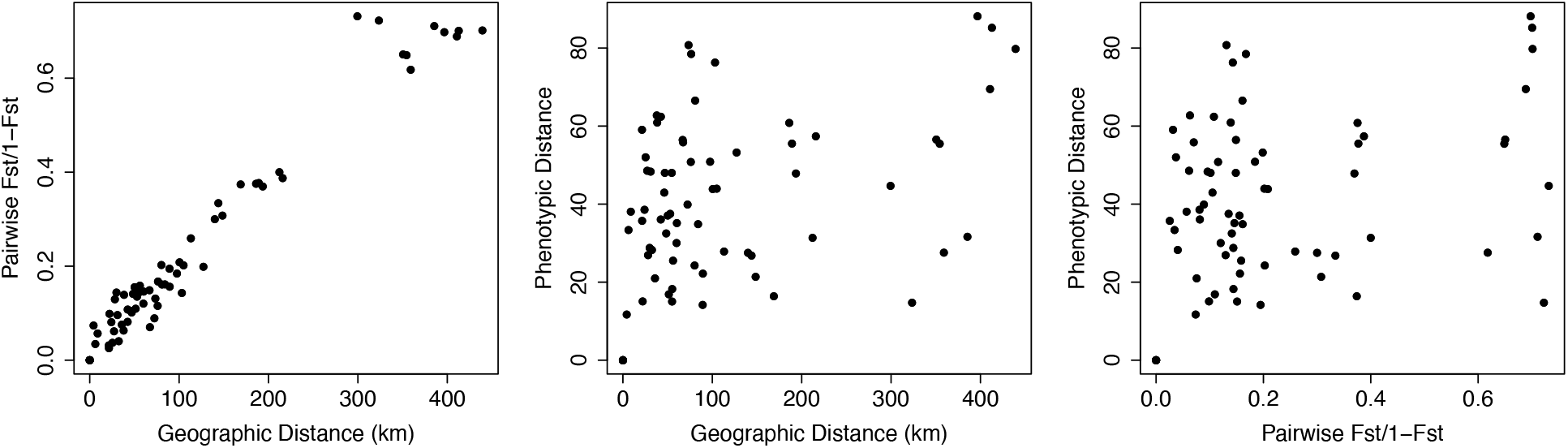
Relationship between over-land geographic distance and genetic distance for 12 sites of red-eyed treefrogs (*Agalychnis callidryas*) along the Pacific coast of Costa Rica. There is a strong positive relationship between geographic and genetic distance, suggesting a pattern of isolation by distance (*r* = 0.965, *p* < 0.001); the relationships between phenotypic and geographic distance (*r* = 0.336, *p* = 0.0085) and genetic and phenotypic distance (*r* = 0.219, *p* = 0.106) were considerably weaker.

We used the R package conStruct to quantify genetic clustering while accounting for geographic distance. We ran conStruct on four sets of individuals: (i) the full dataset, (ii) without the Nicoya Peninsula (Sites 2-12), (iii) northern sites (Site 1-2), and (iv) southern sites (Sites 5-12). Cross-validation analyses showed that spatial models have better predictive accuracy than non-spatial models for all data subsets and *K*-values (Suppl. Fig. 11).

ConStruct results from the full dataset (i) show that non-spatial models inferred more discrete genetic structure compared to spatial models (Fig. 6). The non-spatial model at *K* = 2 shows a separation between the Nicoya Peninsula and other sites, and at *K* = 3 shows a separation between the Osa Peninsula and other sites. Although each additional layer from *K* = 1 to 6 increased the predictive accuracy of the model (Suppl. Fig. 11), layer contributions for the spatial model decreased dramatically after *K* = 1 (Suppl. Fig. 12). The spatial model at *K* = 3 assigned individuals on the Nicoya and Osa Peninsulas similar admixture proportions. There is no obvious biological explanation for why the populations at these sites would be assigned to the same discrete cluster. It is likely that runs at *K* ≥ 2 are describing the genetic dissimilarity between frogs on the Nicoya and Osa Peninsulas with the IBD component of the model within a single layer. This may not reflect recent shared ancestry, and instead may be an artifact of the large geographic distance between these locations combined with gaps in our sampling. Therefore, we interpret *K* = 1 as the model with the most biological significance, and subset the dataset to investigate genetic trends within restricted geographic areas.

**Figure 6.**
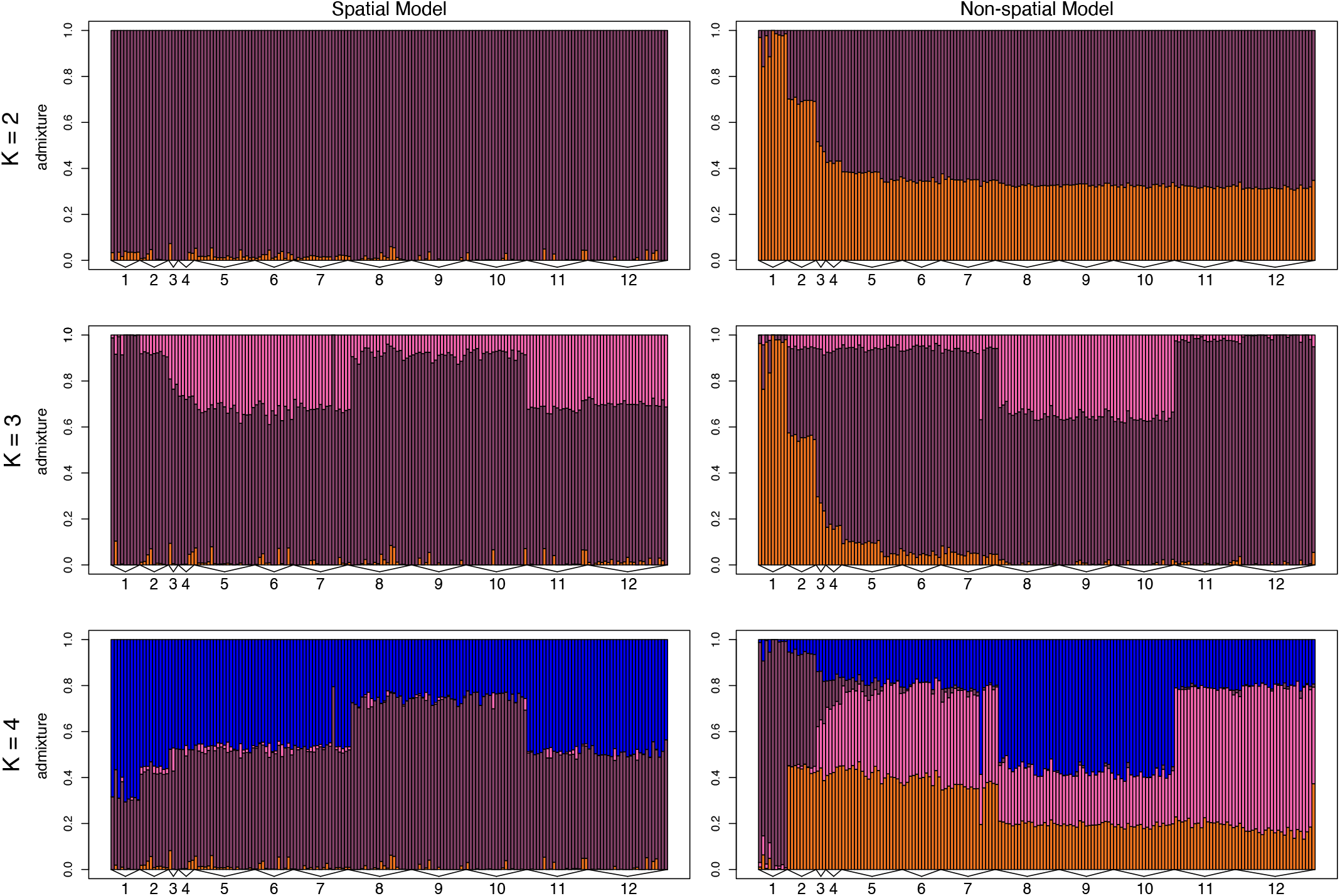
Barplots of admixture proportions for red-eyed treefrogs (*Agalychnis callidryas*) sampled from 12 sites along the Pacific coast of Costa Rica. Admixture proportions shown were generated using conStruct for *K* = 2–4 for both non-spatial and spatial models. Each bar represents an individual. The color of the bar shows the proportion of the individual’s genome that is assigned to each of *K* layers.

We ran conStruct on a subset of the data that excluded the Nicoya Peninsula (ii). Results show very little assignment to the second layer (Fig. 7), and layer contributions are minimal after *K* = 1 (Suppl. Fig. 12). For genetic clustering with just northern sites (iii), there was a difference in assignment between sites, and layer contributions indicate that the most biological relevant *K*-value is *K* = 2 (Fig. 7, Suppl. Fig. 12). When considering the southern sites (iv), individuals on the Osa Peninsula and mainland are partially assigned to different layers, but layer contributions indicate that the most biological relevant *K*-value is *K* = 1 (Fig. 7, Suppl. Fig. 12). Admixture proportions for additional *K*-values and nonspatial models are available in the supplemental material (Suppl. Fig. 13-17). Partitioning the data allowed us to investigate discrete processes within restricted geographic ranges that might not be inferred in the full analysis.

**Figure 7:**
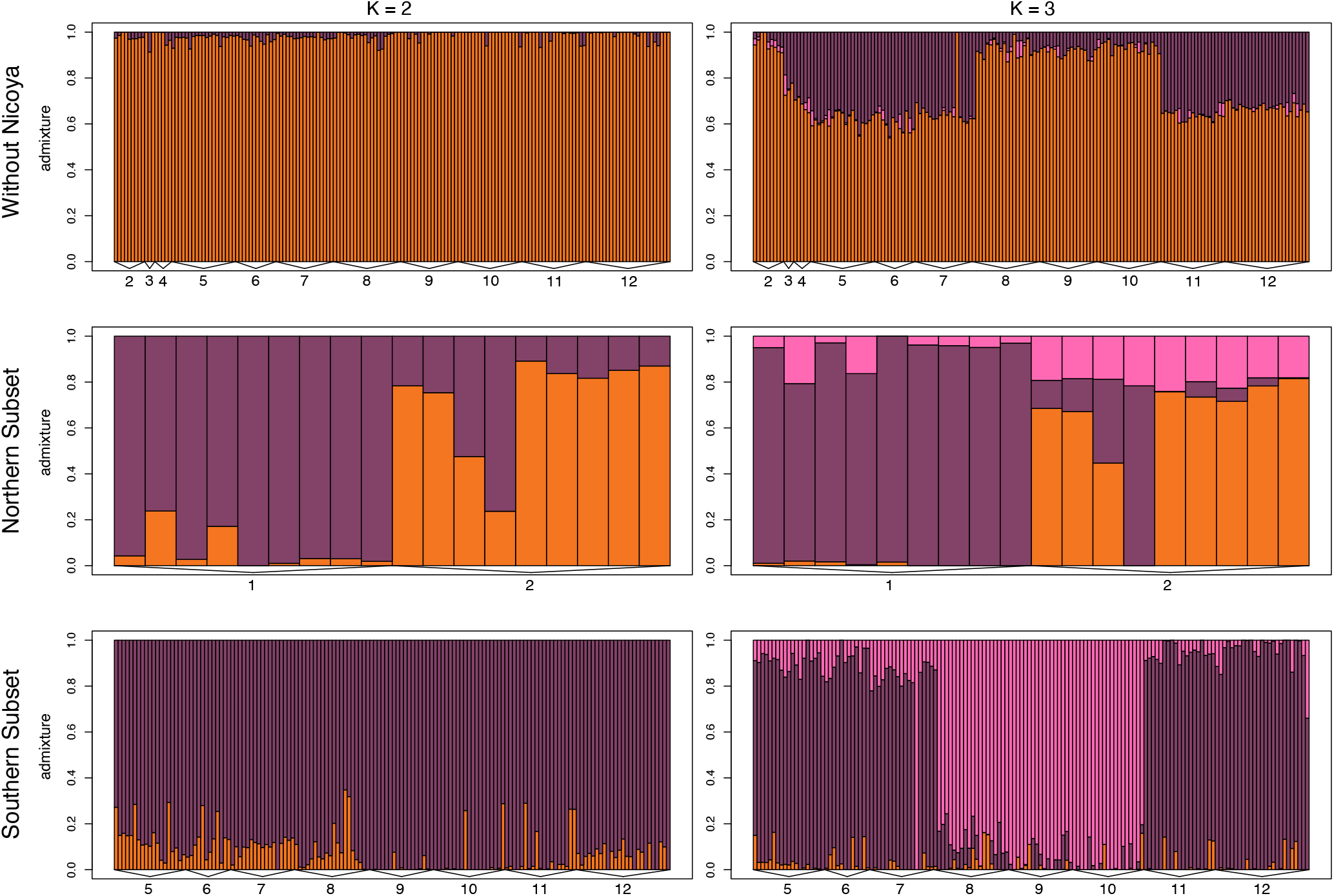
Barplots of admixture proportions for red-eyed treefrogs (*Agalychnis callidryas*) sampled from three data subsets: without Nicoya (Sites 2–12), northern sites (Sites 1–2), and southern sites (5-12). Admixture proportions shown were generated using conStruct for *K* = 2–3 for spatial models. Each bar represents an individual. The color of the bar shows the proportion of the individual’s genome that is assigned to each of K layers.

We used Bayesian linear models to evaluate relationships between phenotypic measurements (color-pattern PC1, hue, saturation, and brightness) and latitude while accounting for genetic covariance. Using 95% credible intervals, with a Dunn-Bonferroni correction for multiple tests, we found that individual leg saturation, hue, and color-pattern PC1, but not brightness, vary significantly with latitude while accounting for genetic covariance among sampled individuals (Fig. 8). These correlations are not significant when using phenotypic averages for each site. We did not find significant relationships between variation in phenotypic measures and θ_w_, or latitude (Suppl. Fig. 18, 19).

**Figure 8.**
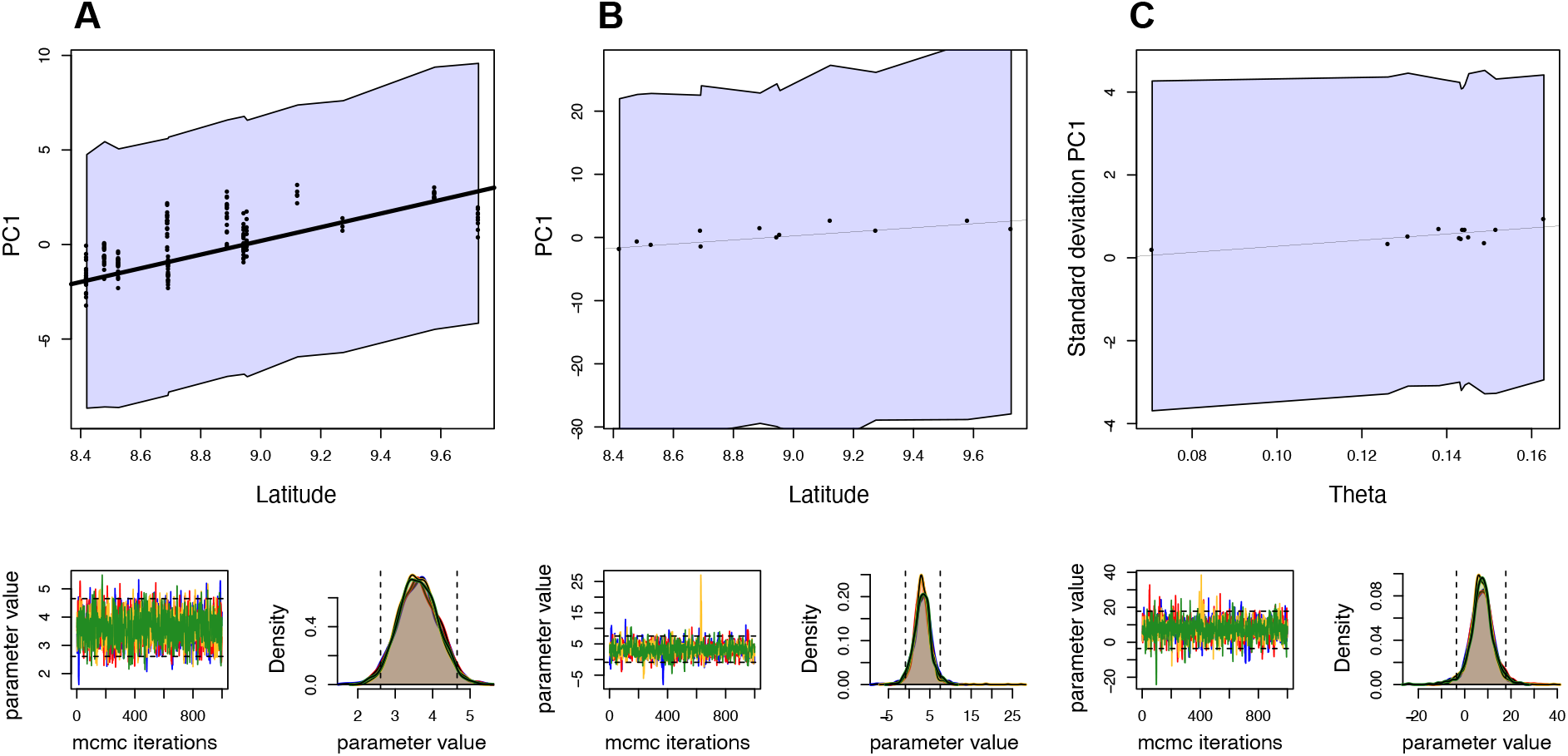
Results of Bayesian models with Dunn-Bonferroni corrected credible intervals (CI) showing the relationships between (A) latitude and color-pattern PC1, (B) latitude and average PC1 value per site, and (C) Wu and Watterson’s theta and variation in PC1 at each site. Each model contains genetic covariance as a covariate. Bolded line indicates that the 95% CI did not overlap with zero. Insets below model fit show estimated parameter values for 5,000 Markov chain Monte Carlo iterations.

## Discussion

The distribution of color pattern in natural systems is complex and can reveal underlying evolutionary dynamics when compared to genetic patterns. Red-eyed treefrogs display dramatic variation in color pattern, yet the two primary morphs found on the Pacific coast of Costa Rica are not restricted to distinct genetic demes (Hypothesis 1B, 3B). Rather, we identified a transition from orange to purple legs and polymorphic sites that contain orange, purple, and novel morphs (Hypothesis 2B) that cannot be explained by recent hybridization between discrete ancestral genetic groups (Hypothesis 4B). Our analyses of the complex relationship between genetic and color-pattern variation along a narrow stretch (∼215 km) of the Pacific coast of Costa Rica suggest that selective forces plus patterns of IBD influence color-pattern distribution in the region.

### Polytypic patterns: Pattern of isolation by distance found across polytypic regions

We found that red-eyed treefrogs at some sites had relatively uniform leg color-pattern, such as orange frogs on the Nicoya Peninsula and purple frogs in the south region, whereas other sites, such as those in the C2 region, were variable (Table 2). Across the Pacific coast, most of the genetic variation among red-eyed treefrog populations is explained by over-land distance among sites; orange and purple morphs do not represent separate genetic demes (Fig. 5, 6). Red-eyed treefrogs are likely semi-continuously distributed, yet, as with most population genetic studies, we sampled at discrete sampling locations; this risks over-estimating the number of discrete genetic groups inferred using clustering models (Bradburd et al., 2018; Frantz et al., 2009; Meirmans, 2012). Therefore, we used conStruct to compare models of population structure with and without spatial information (Bradburd et al., 2018). The nonspatial conStruct models infer more genetic clustering than the spatial models, indicating that clustering in nonspatial models is due to the geographic distance between sites (Fig. 6, 7). Our results highlight the importance of considering patterns of IBD as they relate to genetic relatedness among discrete sampling sites: we would have both over-estimated the number of genetic demes and come to strikingly different conclusions about how diversity is distributed had we relied solely on nonspatial clustering models.

Spatial conStruct models show a division between the mainland and the two peninsular regions (Nicoya and Osa), but we assert that K=1 best describes the data because there is no *a priori* reason to expect that sites on the Nicoya and Osa Peninsulas exchange migrants or share a more recent common ancestor with each other than with neighboring mainland sites. In fact, pairwise *F*_ST_ values between the Nicoya site and Osa sites are among the highest found in the dataset (Table 3, 0.408-0.415). Further, the non-spatial model did not support shared ancestry between Nicoya and Osa.

On both the Nicoya and Osa Peninsulas, we expected red-eyed treefrogs to be isolated from neighboring sites on the mainland: Site 1 at the tip of the Nicoya Peninsula is one of the only known populations of red-eyed treefrogs in the region, as dry forest unsuitable for red-eyed treefrogs cover most of far northwestern Costa Rica, including the Nicoya Peninsula (Kohlmann et al., 2002; Savage, 2002). The Osa Peninsula (Site 8-10) is genetically isolated from mainland sites for many other taxa, including snakes and other frogs (Crawford, 2003; Crawford et al., 2007; Zamudio & Greene, 1997), and we thought this might also be true for red-eyed treefrogs.

Using the full dataset, we did not detect more genetic divergence between mainland and peninsular sites than would be expected given their geographic distance and the rate at which genetic divergence accrues at smaller spatial scales. However, when we restricted analyses to peninsular and nearby mainland sites, we see divergence between the Nicoya Peninsula and mainland, but not between the mainland and the Osa Peninsula (Fig. 7, Suppl. Fig. 12). This suggests that the total divergence between red-eyed treefrogs on the Nicoya Peninsula and mainland is explained by both IBD and barriers to gene flow. As dry forest on the Nicoya Peninsula likely acts as a barrier to gene flow, it is possible that populations along the coast (e.g., Gonzalez, 2013) provide a path for gene flow between the peninsula and mainland. Other studies that have found divergence between the Osa Peninsula and mainland populations have not explicitly incorporated IBD into their analyses, which may explain the discrepancies between our conclusions.

### Polymorphic patterns: Hybridization does not underlie color pattern polymorphism

We tested whether color-pattern polymorphism at a particular site arose through recent hybridization of divergent parent populations, the retention of ancestral polymorphism, and/or ancient secondary contact. We did not find evidence of discrete parental groups, nor recent hybridization in the Central 2 region, despite the color-pattern polymorphism in this region (Table 2, Fig. 2). Rather our findings support that either color-pattern polymorphism was retained from an ancestral population, there has been ancient secondary contact with extended gene flow spreading alleles through the system, and/or that novel morphs have been generated by new mutations. Lab-bred hybrids provided examples of admixed ancestry (F1 stage) and were the result of crossing orange (Site 2) and purple (Site 12) morph parents from geographically distant locations. Thus, these parents were more genetically differentiated than the wild-caught orange and purple morph individuals living in the transition zone. Orange and purple frogs to the north, south, and within the transition zone were assigned to similar genetic groups in both our spatial and nonspatial conStruct analyses, eliminating the possibility of genetic hybrids. Strong signals of IBD (Fig. 5), suggests that gene flow among sites is uninhibited by geographic barriers or other forms of premating reproductive isolation.

While the distribution of color-pattern variation on the Pacific slope of Costa Rica seemingly mirrors patterns on the Caribbean, the underlying processes differ. Red-eyed treefrogs on both the Pacific and Caribbean exhibit color-pattern polymorphism located within a “transition zone” between two distinct morphs. However, polymorphism in the Caribbean is the result of hybridization between two distinct genetic groups of different color morphs (Akopyan et al., 2020b). There, red and blue morph frogs meet in a contact zone and produce frogs with either a novel purple color pattern or a red/blue phenotype distinct from either parental population. Our contrasting findings for the Pacific slope emphasize that diverse evolutionary forces can produce similar patterns of phenotypic variation even within intraspecific lineages. Whereas there is direct evidence that polymorphism on the Caribbean coast is the result of hybridization, the lack of recent hybridization along the Pacific slope suggests that variation in leg phenotype could have arisen due to the retention of ancestral color-pattern polymorphism in the region (as in Corl, Davis, Kuchta, & Sinervo, 2010), or ancient secondary contact (as in Lim et al., 2010). Additionally, polymorphism could be generated by novel mutations (as in Rosenblum et al., 2006, 2010).

### Spatial patterns of phenotype and genotype explained by multiple processes

Patterns of phenotypic and genetic variation permit us to test hypotheses about the evolutionary forces that influence lineage divergence. We found that genetic relatedness predicts phenotypic similarity in some regions of the Pacific coast, but not in others.

Evolution via drift is a common pattern for insular populations (e.g., Knopp et al., 2007, Lehtonen et al., 2009). In our study system, frogs on the Nicoya Peninsula had the highest pairwise *F*_ST_ values to sampled populations at other locations (Table 3), the lowest nucleotide diversity, and highest number of private alleles (Table 1). Individuals from the Nicoya Peninsula have a bright orange phenotype that is distinct from mainland orange frogs (Table 2). Although we cannot rule out local adaptation driving both phenotypic and genetic divergence, red-eyed treefrogs on the Nicoya Peninsula are isolated from the mainland by both geographic distance and tropical dry forests (see above), suggesting that genetic and phenotypic differentiation in this region could have resulted from genetic drift.

In other instances along the Pacific coast, we found that phenotypic similarity is not well predicted by neutral genetic relatedness at our genotyped loci, supporting the hypothesis that color pattern evolves through selection in this region (see below for discussion of selection). Orange and purple morph frogs along the mainland coast and in the polymorphic Central 2 region were not part of separate genetic demes, and instead conformed well to the coast-wide pattern of genetic IBD (Fig. 6,7), despite different phenotypes (Table 2; Fig. 2). We found additional incongruent patterns of variation on the Osa Peninsula, where the three sampled sites are genetically similar (Fig. 3, 6, 7), but phenotypically distinct from one another (Table 2). These incongruences between genetic patterns and phenotype suggest that phenotypic differences are maintained by selection in the presence of gene flow. Whether color-pattern differentiation in these regions was generated by novel mutations, ancestral polymorphisms, and/or the result of genetic drift in these relativity isolated and small populations remains untested. Phenotypic divergence, despite genetic homogeneity, is a common pattern in nature, and usually has been attributed to sexual selection or adaptation to environmental variation. Some examples include *Epipedobates* poison frogs in Peru and Colombia (Tarvin et al., 2017), mimic poison frogs in Peru (Twomey et al., 2013) and redpoll finches in the Arctic circle (Mason & Taylor, 2015). The phenotypic variation on the Osa Peninsula is particularly surprising given the small geographic scale (distance among sites ranges from 20–33 km).

As reported above, the transition in phenotype from orange legs to purple legs is not the result of recent hybridization between two genetic demes meeting at a contact zone. It is possible that it was facilitated by ancient secondary contact between ancestral demes, one with orange flanks and legs and the other with purple flanks and legs, and that color polymorphism is the result of continued gene flow (Endler, 1977). Frogs from the southern sites (7-12) and the Osa Peninsula are part of a different mtDNA clade (clade A, Table 1) from the north (clade B; Table 1; Robertson and Zamudio 2009). This biogeographic discordance between mtDNA and nuclear loci suggests past secondary contact (as in Toews & Brelsford, 2012). Along the mainland Pacific slope, frogs belong to the same genetic cluster, but have different mitochondrial haplotypes and distinct color patterns. This suggests that ancestral mitochondrial groups and color-pattern divergence have been maintained over time, whereas nuclear structure has been lost as a result of gene flow. However, it is also possible that the geographic structure of mtDNA haplotypes is the result of IBD alone (Irwin, 2002). Differentiating between these scenarios would require additional genomic data from these regions. These results highlight the complexity and nuance of this natural system, where multiple processes potentially shape diversity at a fine spatial scale, including genetic drift, selection, and ancient evolutionary history.

Because there is spatial autocorrelation in color pattern along the Pacific coast, it is difficult to decouple phenotypic changes from patterns of IBD to explore possible drivers of color-pattern variation. We therefore used Bayesian linear models to test whether color-pattern phenotype differences are greater than expected given genetic relatedness, and whether those differences are correlated with latitude or genetic diversity. Specifically, we were interested if ecological changes correlated with latitude might play a role in color-pattern variation in red-eyed treefrogs, and if sites with more genetic variation might have more phenotypic variation. We found a significant effect of latitude on phenotype for color-pattern PC1 and leg saturation and hue, but not for brightness measures (Fig. 8, Suppl. Fig. 18, 19). The relationship between phenotype and latitude disappeared at the site level, possibly due to reduced sample size. Future work could investigate the relationship between phenotype and latitude to explore the potential for localized ecological selection on color pattern.

### Role and mechanisms of selection

A complex set of selective pressures likely underlies the genetic and phenotypic patterns observed between and within red-eyed treefrog populations. Although hidden during the day when the frogs are at rest, the flanks and legs of red-eyed treefrogs are visible during activity at night, including during mating aggregations (Pyburn, 1970). Color pattern serves as a species and population-recognition cue in mate choice (Kaiser et al., 2018; Robertson & Greene, 2017). Mate preference for local color pattern has been demonstrated in this species: females from Site 2 (orange) and Site 12 (purple) have a preference for males with their local color pattern in mate-choice trials, suggesting a role for sexual selection on visual signals (Akopyan et al., 2018; Jacobs et al., 2016). Frogs from these sites also have distinguishable male calls and exhibit differences in female courtship behaviors (Akopyan et al., 2018), suggesting that preference for male traits could maintain local color-pattern differences even in the face of gene flow.

Other forms of selection could also shape geographic variation in color pattern. Red-eyed treefrogs secrete noxious host-defense polypeptides from their skin (Conlon et al., 2007; Davis et al., 2016; Mignogna et al., 1997; Sazima, 1974), presenting the possibility that flank- and leg-color patterns act as aposematic signals to visual predators (Robertson & Greene, 2017). Geographic variation in host-defense polypeptides is correlated with color pattern in red-eyed treefrogs, indicating a role for ecological selection (Clark, 2019; Davis et al., 2016). Aposematic coloration is common among other brightly colored anurans (e.g., many species within Dendrobatidae, Mantellidae), and has been shown to vary among and within populations (Klonoski et al., 2019; Maan & Cummings, 2008; Rojas et al., 2014; Roland et al., 2017; Summers & Clough, 2001; Tarvin et al., 2017). Variation in the color pattern of aposematic species is often attributed to the interplay between ecological and sexual selection (Maan & Cummings, 2008; Nokelainen et al., 2012; Rojas & Endler, 2013). Future studies of red-eyed treefrogs that test the role of color pattern in predator avoidance would inform how ecological and sexual selective pressures interact to shape color-pattern distribution.

### Conclusions

Our results highlight the complex evolutionary dynamics that shape color pattern in natural systems. We found that, despite a dramatic and discrete phenotypic change from orange to purple morphs, genetic variation in our study region follows a continuous pattern of IBD. Phenotypic polymorphism at sites located where the orange morph transitions to purple cannot be explained by recent hybridization as the orange and purple morphs are not genetically differentiated groups. Comparison of color pattern and genetic relatedness indicates that selective forces have likely maintained color-pattern divergence in some regions, while genetic drift may play a role in others. Understanding the relative strength of sexual and ecological selection on color pattern is a logical next step for this system.

## Acknowledgements

We thank K. Kaiser and R.E. Espinoza for insightful discussion of the red eyed treefrog system, R. Toczydlowski for guidance with private allele calculation, and Sirena Biological Station and the Firestone Center for Restoration Ecology for access to their properties for sampling. This work was supported by the Department of Biology and Research and Sponsored Projects at California State University, Northridge (CSUN), the CSUN Associated Students Scholarship in Honor of Jolene Koester, and a CSUN tuition waiver (awarded to M.I.C). Research reported in this publication was supported by the National Institute Of General Medical Sciences of the National Institutes of Health under Award Number R35GM137919 (awarded to G.S.B.). This study was conducted in accordance with the current laws of Costa Rica and with approval of the Sistema Nacional de Áreas de Conservación of the Ministerio de Ambiente y Energía of Costa Rica (research and export permits R-053-2015 and 2015-CR1678). All procedures involving animals were approved by the California State University, Northridge Institutional Animal Care and Use Committee (IACUC 1819-005 and 1415-007a).

## Data Accessibility

Raw data, input files, and analysis scripts are available on figshare at https://doi.org/10.6084/m9.figshare.14622744.v2. Bioinformatic scripts available at https://doi.org/10.6084/m9.figshare.11923017.v2.

## Author Contributions

MIC, JMR, MA, EBR and AV designed this study. JMR, MA, and AV carried out field sampling. MIC performed phenotypic analyses, library preparation and bioinformatics. GSB and MIC carried out statistical and computational modeling. Funding support from JMR, EBR, MIC. MIC drafted the manuscript, and all authors have contributed to and approved the final version.

