## Supplemental materials for "Genetic isolation by distance underlies color pattern divergence in red-eyed treefrogs (*Agalychnis callidryas*)"

Supplementary Table 1: Region, site and ID, color pattern measurements (hind leg saturation, brightness, hue, percent gray patch), collection year, and camera used for photography of red-eyed treefrogs (*Agalychnis callidryas*) along the Pacific coast of Costa Rica.

| Region | Site and ID | Saturation | Brightness | Hue | Patch % | Year collected | Camera |
|---|---|---|---|---|---|---|---|
| Nicoya | CAB_951 | 99 | 96 | 28 | 0 | 2004 | Nikon Coolpix 5700 |
| Nicoya | CAB_953 | 99 | 87 | 31 | 0 | 2004 | Nikon Coolpix 5700 |
| Nicoya | CAB_954 | 99 | 80 | 28 | 0 | 2004 | Nikon Coolpix 5700 |
| Nicoya | CAB_955 | 99 | 98 | 31 | 0 | 2004 | Nikon Coolpix 5700 |
| Nicoya | CAB_957 | 99 | 94 | 21 | 0 | 2004 | Nikon Coolpix 5700 |
| Nicoya | CAB_958 | 99 | 89 | 26 | 0 | 2004 | Nikon Coolpix 5700 |
| Nicoya | CAB_959 | 99 | 90 | 29 | 0 | 2004 | Nikon Coolpix 5700 |
| Nicoya | CAB_960 | 99 | 90 | 28 | 0 | 2004 | Nikon Coolpix 5700 |
| Nicoya | CAB_963 | 99 | 84 | 27 | 0 | 2004 | Nikon Coolpix 5700 |
| North | BIJ_010 | 65 | 86 | 25 | 0 | 2015 | Canon PowerShot SX160 |
| North | BIJ_011 | 49 | 67 | 16 | 0 | 2015 | Canon PowerShot SX160 |
| North | BIJ_012 | 90 | 60 | 18 | 0 | 2015 | Canon PowerShot SX160 |
| North | BIJ_05 | 79 | 64 | 29 | 0 | 2015 | Canon PowerShot SX160 |
| North | BIJ_111 | 98 | 47 | 23 | 0 | 2015 | Canon PowerShot SX160 |
| North | BIJ_15 | 87 | 51 | 20 | 0 | 2015 | Canon PowerShot SX160 |
| North | BIJ_16 | 82 | 62 | 33 | 0 | 2015 | Canon PowerShot SX160 |
| North | BIJ_2351 | 99 | 63 | 1 | 0 | 2005 | Nikon Coolpix 5700 |
| North | BIJ_2354 | 99 | 73 | 21 | 0 | 2005 | Nikon Coolpix 5700 |
| C1 | FIR_06 | 66 | 42 | 27 | 0 | 2015 | Canon PowerShot SX160 |
| C1 | FIR_09 | 87 | 55 | 24 | 0 | 2015 | Canon PowerShot SX160 |
| C1 | FIR_14 | 77 | 57 | 19 | 0 | 2015 | Canon PowerShot SX160 |
| C1 | UVI_834 | 99 | 77 | 37 | 0 | 2004 | Nikon Coolpix 5700 |
| C1 | UVI_835 | 98 | 76 | 33 | 0 | 2004 | Nikon Coolpix 5700 |
| C1 | UVI_836 | 98 | 90 | 38 | 20.09 | 2004 | Nikon Coolpix 5700 |
| C1 | UVI_838 | 99 | 66 | 35 | 11.73 | 2004 | Nikon Coolpix 5700 |

| | | | | | | | |
|---|---|---|---|---|---|---|---|
| C1 | UVI_839 | 99 | 89 | 38 | 0 | 2004 | Nikon Coolpix 5700 |
| C2 | CRT_364 | 71 | 33 | 16 | 9.54 | 2015 | Canon PowerShot SX160 |
| C2 | CRT_365 | 88 | 61 | 28 | 0 | 2015 | Canon PowerShot SX160 |
| C2 | CRT_366 | 77 | 29 | 22 | 15.17 | 2015 | Canon PowerShot SX160 |
| C2 | CRT_367 | 83 | 48 | 15 | 0 | 2015 | Canon PowerShot SX160 |
| C2 | CRT_368 | 85 | 32 | 20 | 35.21 | 2015 | Canon PowerShot SX160 |
| C2 | CRT_369 | 85 | 65 | 29 | 17.48 | 2015 | Canon PowerShot SX160 |
| C2 | CRT_370 | 81 | 33 | 20 | 0 | 2015 | Canon PowerShot SX160 |
| C2 | CRT_372 | 78 | 25 | 16 | 0 | 2015 | Canon PowerShot SX160 |
| C2 | CRT_373 | 80 | 29 | 15 | 0 | 2015 | Canon PowerShot SX160 |
| C2 | CRT_374 | 80 | 48 | 23 | 4.8 | 2015 | Canon PowerShot SX160 |
| C2 | CRT_375 | 74 | 22 | 20 | 29.44 | 2015 | Canon PowerShot SX160 |
| C2 | CRT_376 | 75 | 31 | 16 | 0 | 2015 | Canon PowerShot SX160 |
| C2 | PSR_377 | 71 | 28 | 14 | 32.29 | 2015 | Canon PowerShot SX160 |
| C2 | PSR_378 | 74 | 34 | 18 | 9.02 | 2015 | Canon PowerShot SX160 |
| C2 | PSR_379 | 64 | 79 | 26 | 20.99 | 2015 | Canon PowerShot SX160 |
| C2 | PSR_380 | 85 | 48 | 16 | 10.08 | 2015 | Canon PowerShot SX160 |
| C2 | PSR_381 | 88 | 79 | 23 | 9.67 | 2015 | Canon PowerShot SX160 |
| C2 | PSR_382 | 58 | 35 | 14 | 18.12 | 2015 | Canon PowerShot SX160 |
| C2 | PSR_383 | 71 | 31 | 15 | 15.55 | 2015 | Canon PowerShot SX160 |
| C2 | PSR_384 | 61 | 63 | 16 | 26.1 | 2015 | Canon PowerShot SX160 |
| C2 | PSR_385 | 79 | 36 | 20 | 10.26 | 2015 | Canon PowerShot SX160 |
| C2 | PSR_386 | 69 | 33 | 11 | 4.08 | 2015 | Canon PowerShot SX160 |
| C2 | PSR_387 | 82 | 71 | 24 | 20.66 | 2015 | Canon PowerShot SX160 |
| C2 | PSR_388 | 70 | 30 | 8 | 16.52 | 2015 | Canon PowerShot SX160 |
| C2 | PSR_389 | 72 | 35 | 15 | 26.03 | 2015 | Canon PowerShot SX160 |
| C2 | PSR_390 | 79 | 40 | 22 | 0 | 2015 | Canon PowerShot SX160 |
| C2 | PSR_391 | 77 | 30 | 14 | 13.77 | 2015 | Canon PowerShot SX160 |
| C2 | PSR_392 | 70 | 33 | 12 | 0 | 2015 | Canon PowerShot SX160 |

| | | | | | | | |
|---|---|---|---|---|---|---|---|
| C2 | PSR_393 | 67 | 39 | 20 | 25.24 | 2015 | Canon PowerShot SX160 |
| C2 | PSR_394 | 75 | 55 | 20 | 16.65 | 2015 | Canon PowerShot SX160 |
| C2 | PSR_395 | 72 | 57 | 22 | 21.31 | 2015 | Canon PowerShot SX160 |
| C2 | PSR_396 | 70 | 36 | 15 | 9.94 | 2015 | Canon PowerShot SX160 |
| C2 | SIE_742 | 80 | 66 | 26 | 36.06 | 2004 | Nikon Coolpix 5700 |
| C2 | SIE_745 | 66 | 49 | 33 | 47.4085653 | 2004 | Nikon Coolpix 5700 |
| C2 | SIE_748 | 90 | 82 | 38 | 26.13 | 2004 | Nikon Coolpix 5700 |
| C2 | SIE_749 | 73 | 65 | 23 | 21.6 | 2004 | Nikon Coolpix 5700 |
| C2 | SIE_750 | 99 | 77 | 23 | 0 | 2004 | Nikon Coolpix 5700 |
| C2 | SIE_751 | 99 | 80 | 27 | 28.57 | 2004 | Nikon Coolpix 5700 |
| C2 | SIE_752 | 80 | 85 | 27 | 21.48 | 2004 | Nikon Coolpix 5700 |
| C2 | SIE_753 | 92 | 98 | 33 | 5.61 | 2004 | Nikon Coolpix 5700 |
| C2 | SIE_754 | 67 | 75 | 25 | 0 | 2004 | Nikon Coolpix 5700 |
| C2 | SIE_755 | 99 | 79 | 29 | 0 | 2004 | Nikon Coolpix 5700 |
| C2 | SIE_756 | 78 | 67 | 29 | 24.7 | 2004 | Nikon Coolpix 5700 |
| C2 | SIE_757 | 89 | 75 | 24 | 0 | 2004 | Nikon Coolpix 5700 |
| C2 | SIE_758 | 99 | 67 | 24 | 0 | 2004 | Nikon Coolpix 5700 |
| C2 | SIE_759 | 83 | 99 | 29 | 0 | 2004 | Nikon Coolpix 5700 |
| C2 | SIE_780 | 96 | 44 | 13 | 37.15 | 2004 | Nikon Coolpix 5700 |
| C2 | SIE_781 | 46 | 76 | 20 | 24.65 | 2004 | Nikon Coolpix 5700 |
| C2 | SIE_782 | 99 | 78 | 34 | 23.28 | 2004 | Nikon Coolpix 5700 |
| Osa | CAM_809 | 87 | 44 | 18 | 43.46 | 2004 | Nikon Coolpix 5700 |
| Osa | CAM_810 | 99 | 55 | 9 | 21.15 | 2004 | Nikon Coolpix 5700 |
| Osa | CAM_812 | 62 | 64 | 30 | 27 | 2004 | Nikon Coolpix 5700 |
| Osa | CAM_813 | 59 | 55 | 25 | 34.68 | 2004 | Nikon Coolpix 5700 |
| Osa | CAM_814 | 95 | 82 | 30 | 37.38 | 2004 | Nikon Coolpix 5700 |
| Osa | CAM_815 | 99 | 83 | 21 | 0 | 2004 | Nikon Coolpix 5700 |
| Osa | CAM_816 | 99 | 54 | 19 | 0 | 2004 | Nikon Coolpix 5700 |
| Osa | CAM_817 | 96 | 71 | 26 | 0 | 2004 | Nikon Coolpix 5700 |

| | | | | | | | |
|---|---|---|---|---|---|---|---|
| Osa | CAM_818 | 99 | 76 | 5 | 0 | 2004 | Nikon Coolpix 5700 |
| Osa | CAM_819 | 94 | 54 | 18 | 0 | 2004 | Nikon Coolpix 5700 |
| Osa | CAM_820 | 84 | 58 | 30 | 0 | 2004 | Nikon Coolpix 5700 |
| Osa | CAM_821 | 96 | 55 | 14 | 0 | 2004 | Nikon Coolpix 5700 |
| Osa | CAM_822 | 98 | 72 | 14 | 0 | 2004 | Nikon Coolpix 5700 |
| Osa | CAM_823 | 99 | 61 | 12 | 0 | 2004 | Nikon Coolpix 5700 |
| Osa | CAM_824 | 99 | 48 | 10 | 0 | 2004 | Nikon Coolpix 5700 |
| Osa | CAM_825 | 97 | 78 | 22 | 36.15 | 2004 | Nikon Coolpix 5700 |
| Osa | CAM_826 | 91 | 76 | 22 | 39.48 | 2004 | Nikon Coolpix 5700 |
| Osa | SIR_1 | 49 | 71 | 18 | 20.21 | 2015 | Canon PowerShot SX160 |
| Osa | SIR_11 | 51 | 55 | 14 | 29.07 | 2015 | Canon PowerShot SX160 |
| Osa | SIR_12 | 55 | 57 | 11 | 11.73 | 2015 | Canon PowerShot SX160 |
| Osa | SIR_13 | 34 | 56 | 0 | 29.26 | 2015 | Canon PowerShot SX160 |
| Osa | SIR_14 | 45 | 56 | 12 | 0 | 2015 | Canon PowerShot SX160 |
| Osa | SIR_15 | 38 | 63 | 10 | 13.05 | 2015 | Canon PowerShot SX160 |
| Osa | SIR_16 | 40 | 60 | 21 | 31.71 | 2015 | Canon PowerShot SX160 |
| Osa | SIR_17 | 35 | 75 | 21 | 14.33 | 2015 | Canon PowerShot SX160 |
| Osa | SIR_18 | 24 | 73 | 19 | 26.1 | 2015 | Canon PowerShot SX160 |
| Osa | SIR_19 | 23 | 65 | 11 | 30.08 | 2015 | Canon PowerShot SX160 |
| Osa | SIR_2 | 41 | 56 | 2 | 0 | 2015 | Canon PowerShot SX160 |
| Osa | SIR_20 | 26 | 67 | 23 | 11.63 | 2015 | Canon PowerShot SX160 |
| Osa | SIR_3 | 51 | 60 | 9 | 0 | 2015 | Canon PowerShot SX160 |
| Osa | SIR_4 | 50 | 50 | 7 | 5.3 | 2015 | Canon PowerShot SX160 |
| Osa | SIR_5 | 34 | 70 | 21 | 48.93 | 2015 | Canon PowerShot SX160 |
| Osa | SIR_6 | 51 | 64 | 17 | 15.84 | 2015 | Canon PowerShot SX160 |
| Osa | SIR_7 | 37 | 57 | 13 | 11.7324576 | 2015 | Canon PowerShot SX160 |
| Osa | SIR_8 | 35 | 70 | 18 | 47.7 | 2015 | Canon PowerShot SX160 |
| Osa | SIR_9 | 40 | 54 | 12 | 20.9619935 | 2015 | Canon PowerShot SX160 |
| Osa | TGR_397 | 45 | 36 | 19 | 19.85 | 2015 | Canon PowerShot SX160 |

| | | | | | | | |
|---|---|---|---|---|---|---|---|
| Osa | TGR_398 | 51 | 22 | 6 | 0 | 2015 | Canon PowerShot SX160 |
| Osa | TGR_399 | 49 | 18 | 11 | 29.06 | 2015 | Canon PowerShot SX160 |
| Osa | TGR_400 | 46 | 21 | 10 | 0 | 2015 | Canon PowerShot SX160 |
| Osa | TGR_401 | 50 | 24 | 12 | 0 | 2015 | Canon PowerShot SX160 |
| Osa | TGR_402 | 44 | 19 | 9 | 23.41 | 2015 | Canon PowerShot SX160 |
| Osa | TGR_403 | 48 | 20 | 14 | 0 | 2015 | Canon PowerShot SX160 |
| Osa | TGR_404 | 52 | 22 | 20 | 0 | 2015 | Canon PowerShot SX160 |
| Osa | TGR_405 | 46 | 22 | 14 | 27.5767099 | 2015 | Canon PowerShot SX160 |
| Osa | TGR_406 | 35 | 31 | 16 | 56.8621396 | 2015 | Canon PowerShot SX160 |
| Osa | TGR_407 | 42 | 28 | 22 | 26.651734 | 2015 | Canon PowerShot SX160 |
| Osa | TGR_408 | 48 | 33 | 10 | 16.44 | 2015 | Canon PowerShot SX160 |
| Osa | TGR_409 | 42 | 22 | 13 | 0 | 2015 | Canon PowerShot SX160 |
| Osa | TGR_410 | 46 | 28 | 15 | 30.2 | 2015 | Canon PowerShot SX160 |
| Osa | TGR_411 | 51 | 42 | 13 | 23.02 | 2015 | Canon PowerShot SX160 |
| Osa | TGR_412 | 49 | 27 | 18 | 19.24 | 2015 | Canon PowerShot SX160 |
| Osa | TGR_413 | 37 | 29 | 30 | 23.82 | 2015 | Canon PowerShot SX160 |
| Osa | TGR_414 | 36 | 26 | 17 | 26.78 | 2015 | Canon PowerShot SX160 |
| Osa | TGR_416 | 50 | 41 | 11 | 0 | 2015 | Canon PowerShot SX160 |
| South | GMB_344 | 39 | 38 | 17 | 19.98 | 2015 | Canon PowerShot SX160 |
| South | GMB_345 | 35 | 45 | 24 | 20.47 | 2015 | Canon PowerShot SX160 |
| South | GMB_347 | 36 | 22 | 27 | 59.76 | 2015 | Canon PowerShot SX160 |
| South | GMB_348 | 45 | 18 | 11 | 45.71 | 2015 | Canon PowerShot SX160 |
| South | GMB_349 | 34 | 21 | 20 | 39.23 | 2015 | Canon PowerShot SX160 |
| South | GMB_350 | 41 | 27 | 19 | 49.29 | 2015 | Canon PowerShot SX160 |
| South | GMB_351 | 47 | 30 | 13 | 34.28 | 2015 | Canon PowerShot SX160 |
| South | GMB_352 | 40 | 16 | 11 | 32.81 | 2015 | Canon PowerShot SX160 |
| South | GMB_353 | 44 | 34 | 16 | 45.26 | 2015 | Canon PowerShot SX160 |
| South | GMB_354 | 48 | 26 | 11 | 28.42 | 2015 | Canon PowerShot SX160 |
| South | GMB_355 | 42 | 25 | 16 | 31.58 | 2015 | Canon PowerShot SX160 |

| | | | | | | | |
|---|---|---|---|---|---|---|---|
| South | GMB_356 | 47 | 37 | 18 | 32.87 | 2015 | Canon PowerShot SX160 |
| South | GMB_357 | 47 | 25 | 12 | 30.79 | 2015 | Canon PowerShot SX160 |
| South | GMB_358 | 48 | 28 | 14 | 11.6 | 2015 | Canon PowerShot SX160 |
| South | GMB_359 | 55 | 30 | 11 | 34.85 | 2015 | Canon PowerShot SX160 |
| South | GMB_360 | 51 | 25 | 13 | 13.57 | 2015 | Canon PowerShot SX160 |
| South | GMB_361 | 38 | 39 | 13 | 22.56 | 2015 | Canon PowerShot SX160 |
| South | GMB_362 | 53 | 42 | 13 | 35.22 | 2015 | Canon PowerShot SX160 |
| South | GMB_363 | 54 | 33 | 13 | 14.75 | 2015 | Canon PowerShot SX160 |
| South | PAV_04 | 42 | 43 | -12 | 57.14 | 2015 | Canon PowerShot SX160 |
| South | PAV_06 | 52 | 45 | -14 | 16.69 | 2015 | Canon PowerShot SX160 |
| South | PAV_07 | 43 | 45 | -11 | 10.44 | 2015 | Canon PowerShot SX160 |
| South | PAV_11 | 19 | 48 | -3 | 32.9338868 | 2015 | Canon PowerShot SX160 |
| South | PAV_113 | 21 | 46 | 15 | 37.7190902 | 2015 | Canon PowerShot SX160 |
| South | PAV_13 | 32 | 31 | 7 | 29.5767458 | 2015 | Canon PowerShot SX160 |
| South | PAV_15 | 25 | 58 | 11 | 47.9586752 | 2015 | Canon PowerShot SX160 |
| South | PAV_2182 | 43 | 59 | -1 | 33.05 | 2005 | Canon PowerShot SX160 |
| South | PAV_2183 | 61 | 36 | -6 | 10.84 | 2005 | Canon PowerShot SX160 |
| South | PAV_2184 | 63 | 46 | -14 | 28.01 | 2005 | Canon PowerShot SX160 |
| South | PAV_2185 | 35 | 55 | -7 | 47.78 | 2005 | Canon PowerShot SX160 |
| South | PAV_2186 | 54 | 45 | -4 | 68.75 | 2005 | Canon PowerShot SX160 |
| South | PAV_2189 | 52 | 62 | 4 | 56.79 | 2005 | Canon PowerShot SX160 |
| South | PAV_2190 | 55 | 64 | -6 | 42.24 | 2005 | Canon PowerShot SX160 |
| South | PAV_2191 | 41 | 56 | 6 | 36.44 | 2005 | Canon PowerShot SX160 |
| South | PAV_2192 | 42 | 61 | 4 | 49.34 | 2005 | Canon PowerShot SX160 |
| South | PAV_2193 | 53 | 52 | -3 | 49.71 | 2005 | Canon PowerShot SX160 |
| South | PAV_870 | 41 | 58 | 6 | 58.6509321 | 2004 | Nikon Coolpix 5700 |
| South | PAV_871 | 60 | 52 | 10 | 18.8930818 | 2004 | Nikon Coolpix 5700 |
| South | PAV_872 | 39 | 55 | 16 | 25.3525021 | 2004 | Nikon Coolpix 5700 |
| South | PAV_873 | 35 | 62 | 2 | 32.516507 | 2004 | Nikon Coolpix 5700 |

| | | | | | | | |
|---|---|---|---|---|---|---|---|
| South | PAV_874 | 42 | 60 | 4 | 27.68757 | 2004 | Nikon Coolpix 5700 |
| South | PAV_879 | 40 | 70 | 3 | 9.03069612 | 2004 | Nikon Coolpix 5700 |
| South | PAV_880 | 44 | 56 | -4 | 34.1522075 | 2004 | Nikon Coolpix 5700 |
| South | PAV_883 | 49 | 68 | 8 | 1.5121508 | 2004 | Nikon Coolpix 5700 |

Supplementary Table 2. Discriminant function analysis of leg color saturation values for 12 sampling sites of red-eyed treefrogs (*Agalychnis callidryas*) along the Pacific coast of Costa Rica. Each individual was classified based on leg color saturation and assigned to a sampling site.

| | Region of Origin | | | | | | | | | | | |
|---|---|---|---|---|---|---|---|---|---|---|---|---|
| | Nicoya | North | Central 1 | | Central 2 | | | Osa | | | South | |
| Classification | 1 | 2 | 3 | 4 | 5 | 6 | 7 | 8 | 9 | 10 | 11 | 12 |
| 1 | 0 | 0 | 0 | 0 | 0 | 0 | 0 | 0 | 0 | 0 | 0 | 0 |
| 2 | 0 | 0 | 0 | 0 | 0 | 0 | 0 | 0 | 0 | 0 | 0 | 0 |
| 3 | 0 | 0 | 0 | 0 | 0 | 0 | 0 | 0 | 0 | 0 | 0 | 0 |
| 4 | 0 | 0 | 0 | 0 | 0 | 0 | 0 | 0 | 0 | 0 | 0 | 0 |
| 5 | 0 | 0 | 0 | 0 | 0 | 0 | 0 | 0 | 0 | 0 | 0 | 0 |
| 6 | 0 | 22% | 67% | 0% | 42% | 80% | 24% | 12% | 0 | 0 | 0 | 12% |
| 7 | 0 | 22% | 33% | 0% | 50% | 10% | 18% | 12% | 0 | 0 | 0 | 0 |
| 8 | 100% | 44% | 0 | 100% | 8% | 5% | 53% | 76% | 0 | 0 | 0 | 0 |
| 9 | 0 | 0 | 0 | 0 | 0 | 0 | 0 | 0 | 16% | 0 | 0 | 16% |
| 10 | 0 | 0 | 0 | 0 | 0 | 5% | 0 | 0 | 0 | 0 | 0 | 0 |
| 11 | 0 | 0 | 0 | 0 | 0 | 0 | 0 | 0 | 0 | 0 | 0 | 0 |
| 12 | 0 | 11% | 0 | 0 | 0 | 0 | 6% | 0 | 84% | 100% | 100% | 72% |

Supplementary Table 3. Discriminant function analysis of leg color hue values for 12 sampling sites of red-eyed treefrogs (*Agalychnis callidryas*) along the Pacific coast of Costa Rica. Each individual was classified based on leg color hue and assigned to a sampling site.

| | | Region of Origin | | | | | | | | | | | |
|---|---|---|---|---|---|---|---|---|---|---|---|---|---|
| | | Nicoya | North | Central 1 | | Central 2 | | | Osa | | | South | |
| | | 1 | 2 | 3 | 4 | 5 | 6 | 7 | 8 | 9 | 10 | 11 | 12 |
| Classification | 1 | 0 | 0 | 0 | 0 | 0 | 0 | 0 | 0 | 0 | 0 | 0 | 0 |
| | 2 | 0 | 0 | 0 | 0 | 0 | 0 | 0 | 0 | 0 | 0 | 0 | 0 |
| | 3 | 0 | 0 | 0 | 0 | 0 | 0 | 0 | 0 | 0 | 0 | 0 | 0 |
| | 4 | 0 | 0 | 0 | 60% | 0 | 0 | 6% | 0 | 0 | 0 | 0 | 0 |
| | 5 | 0 | 0 | 0 | 0 | 0 | 0 | 0 | 0 | 0 | 0 | 0 | 0 |
| | 6 | 11% | 44% | 33% | 0 | 50% | 30% | 6% | 24% | 37% | 26% | 32% | 4% |
| | 7 | 89% | 33% | 67% | 40% | 17% | 10% | 82% | 29% | 5% | 5% | 11% | 0% |
| | 8 | 0 | 11% | 0 | 0 | 17% | 15% | 0% | 12% | 0 | 5% | 0 | 0 |
| | 9 | 0 | 0 | 0 | 0 | 0 | 30% | 6% | 29% | 42% | 53% | 58% | 12% |
| | 10 | 0 | 0 | 0 | 0 | 17% | 15% | 0 | 0 | 0 | 5% | 0% | 4% |
| | 11 | 0 | 0 | 0 | 0 | 0 | 0 | 0 | 0 | 0 | 0 | 0 | 0 |
| | 12 | 0 | 11% | 0 | 0 | 0 | 0 | 0 | 6% | 16% | 5% | 0 | 80% |

Supplementary Table 4. Discriminant function analysis of leg color brightness values for 12 sampling sites of red-eyed treefrogs (*Agalychnis callidryas*) along the Pacific coast of Costa Rica. Each individual was classified based on leg color brightness and assigned to a sampling site.

| | Region of Origin | | | | | | | | | | | |
|---|---|---|---|---|---|---|---|---|---|---|---|---|
| | Nicoya | North | Central 1 | | | Central 2 | | | Osa | | South | |
| | 67% | 0 | 0 | 40% | 0 | 0 | 12% | 0 | 0 | 0 | 0 | 0 |
| Classification 1 | 0 | 0 | 0 | 0 | 0 | 0 | 0 | 0 | 0 | 0 | 0 | 0 |
| 2 | 0 | 0 | 0 | 0 | 0 | 0 | 0 | 0 | 0 | 0 | 0 | 0 |
| 3 | 0 | 0 | 0 | 0 | 0 | 0 | 0 | 0 | 0 | 0 | 0 | 0 |
| 4 | 0 | 0 | 0 | 0 | 0 | 0 | 0 | 0 | 0 | 0 | 0 | 0 |
| 5 | 0 | 0 | 33% | 0 | 0 | 10% | 6% | 6% | 0 | 11% | 26% | 16% |
| 6 | 33% | 22% | 0 | 40% | 0 | 15% | 53% | 41% | 26% | 0 | 0 | 4% |
| 7 | 0 | 0 | 0 | 0 | 0 | 0 | 0 | 0 | 0 | 0 | 0 | 0 |
| 8 | 0 | 44% | 0 | 20% | 8% | 5% | 24% | 6% | 21% | 0 | 0 | 16% |
| 9 | 0 | 0 | 0 | 0 | 17% | 5% | 0 | 0 | 0 | 68% | 53% | 0 |
| 10 | 0 | 0 | 0 | 0 | 50% | 50% | 0 | 0 | 0 | 21% | 21% | 8% |
| 11 | 0 | 33% | 67% | 0 | 25% | 15% | 6% | 47% | 53% | 0% | 0% | 56% |
| 12 | 67% | 0 | 0 | 40% | 0 | 0 | 12% | 0 | 0 | 0 | 0 | 0 |

Supplementary Table 5. Discriminant function analysis of leg color patch percentage values for 12 sampling sites of red-eyed treefrogs (*Agalychnis callidryas*) along the Pacific coast of Costa Rica. Each individual was classified based on leg color patch percentage and assigned to a sampling site.

| | Region of Origin | | | | | | | | | | | |
|---|---|---|---|---|---|---|---|---|---|---|---|---|
| | Nicoya | North | Central 1 | | Central 2 | | | Osa | | | South | |
| Classification | 1 | 2 | 3 | 4 | 5 | 6 | 7 | 8 | 9 | 10 | 11 | 12 |
| 1 | 0 | 0 | 0 | 0 | 0 | 0 | 0 | 0 | 0 | 0 | 0 | 0 |
| 2 | 0 | 0 | 0 | 0 | 0 | 0 | 0 | 0 | 0 | 0 | 0 | 0 |
| 3 | 0 | 0 | 0 | 0 | 0 | 0 | 0 | 0 | 0 | 0 | 0 | 0 |
| 4 | 0 | 0 | 0 | 0 | 0 | 0 | 0 | 0 | 0 | 0 | 0 | 0 |
| 5 | 0 | 0 | 0 | 0 | 0 | 0 | 0 | 0 | 0 | 0 | 0 | 0 |
| 6 | 100% | 100% | 100% | 80% | 83% | 65% | 41% | 59% | 53% | 47% | 16% | 24% |
| 7 | 0 | 0 | 0 | 0 | 0 | 0 | 0 | 0 | 0 | 0 | 0 | 0 |
| 8 | 0 | 0 | 0 | 0 | 0 | 0 | 0 | 0 | 0 | 0 | 0 | 0 |
| 9 | 0 | 0 | 0 | 20% | 0 | 15% | 18% | 6% | 11% | 16% | 16% | 0 |
| 10 | 0 | 0 | 0 | 0 | 0 | 0 | 0 | 0 | 0 | 0 | 0 | 0 |
| 11 | 0 | 0 | 0 | 0 | 0 | 0 | 0 | 0 | 0 | 0 | 0 | 0 |
| 12 | 0 | 0 | 0 | 0 | 17% | 20% | 41% | 35% | 37% | 37% | 68% | 76% |

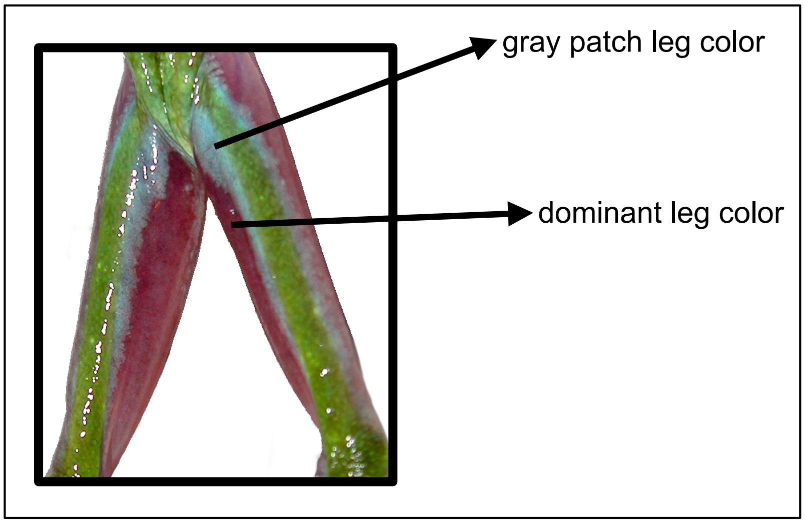


Supplemental Figure 1: Dorsal view of the legs of a red-eyed treefrog (*Agalychnis callidryas*) from Site 12, showing both the purple dominant leg color and the inner blue/gray patch.

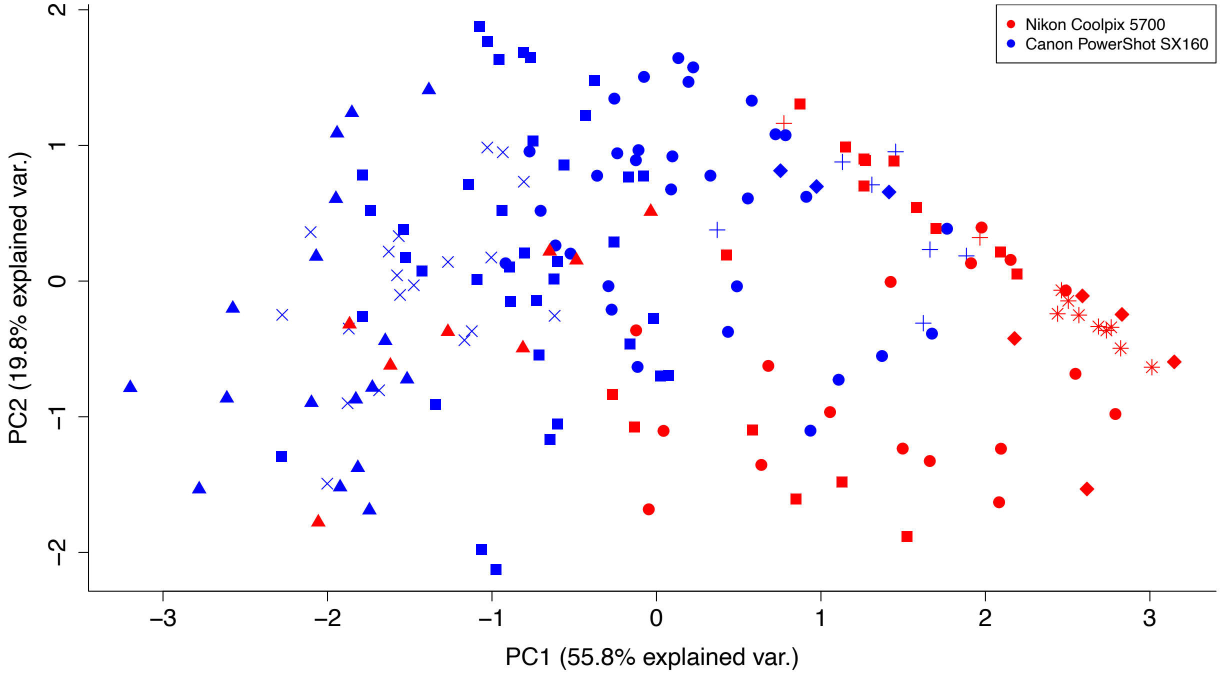


Supplemental Figure 2: Principal component analysis of leg phenotype for red-eyed treefrogs (*Agalychnis callidryas*) sampled from 12 sites along the Pacific coast of Costa Rica. Color indicates camera used for photography. Frogs sampled in 2015 were photographed with a Canon PowerShot SX160. Frogs sampled before 2015 were photographed with a Nikon Coolpix 5700. Symbols correspond to sampling sites.

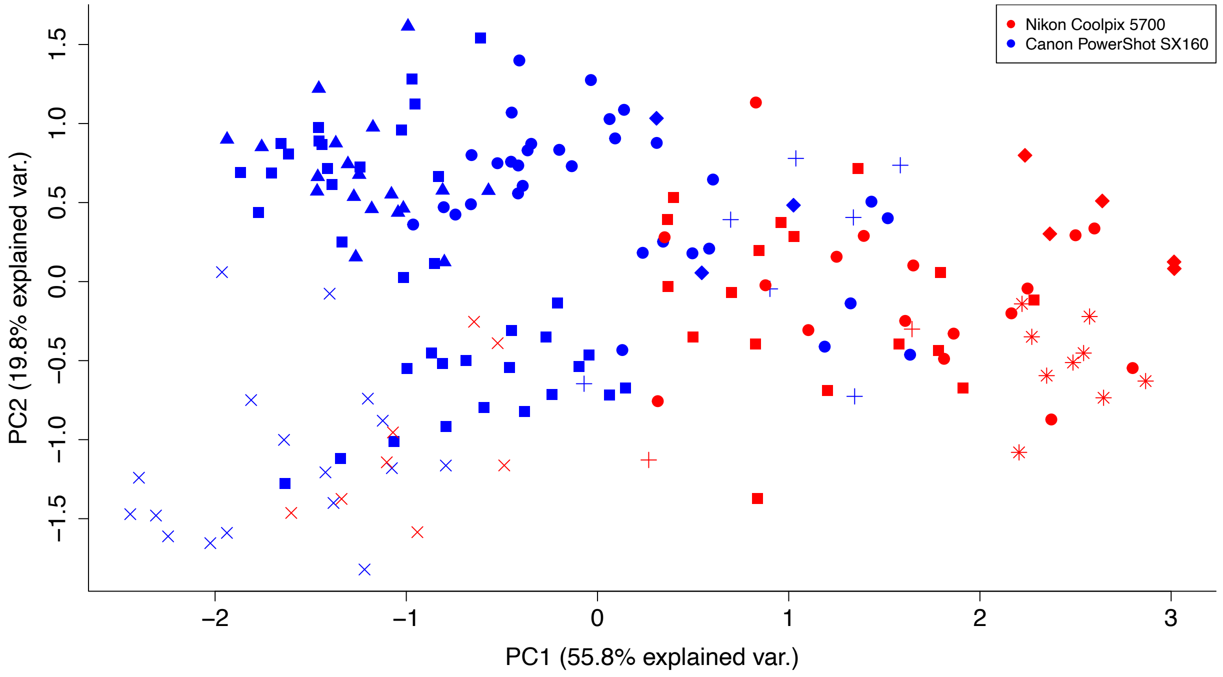


Supplemental Figure 3: Principal component analysis of leg coloration measurements (saturation, brightness, and hue) for red-eyed treefrogs (*Agalychnis callidryas*) sampled from 12 sites along the Pacific coast of Costa Rica. Color indicates sampling year. Frogs sampled in 2015 were photographed with a Canon PowerShot SX160. Frogs sampled before 2015 were photographed with a Nikon Coolpix 5700. Symbols "x" (Site 12) and "+" (Site 2) are frogs from sites that were sampled with both cameras.

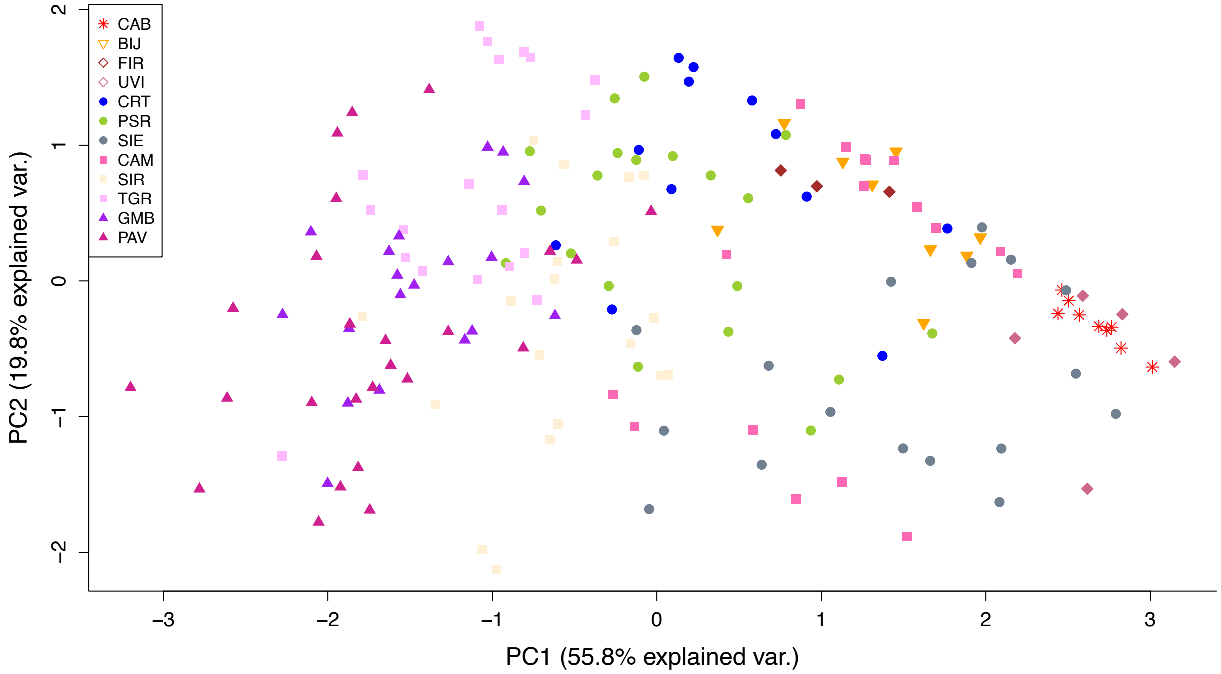


Supplemental Figure 4: Principal component analysis of leg phenotype for red-eyed treefrogs (*Agalychnis callidryas*) sampled from 12 sites along the Pacific coast of Costa Rica. Symbols represent individuals. Symbol type represents region, while color represents sampling site.

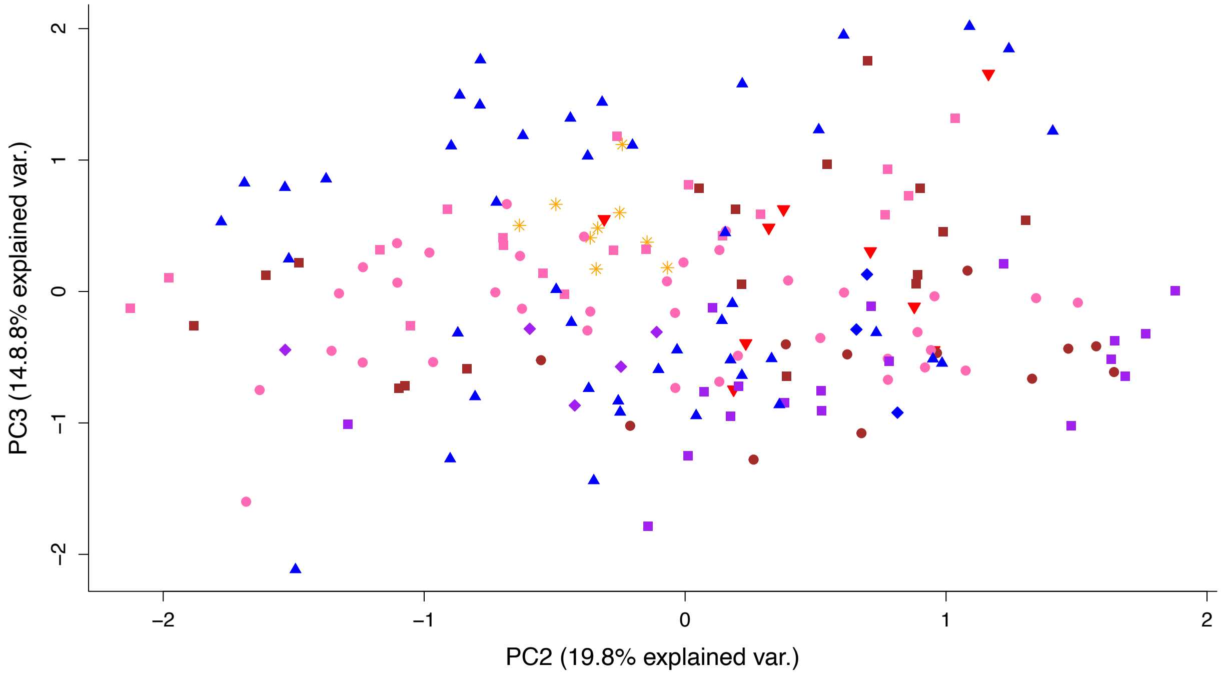


Supplemental Figure 5: Plot of principal components 2 and 3 from principal component analysis of leg color pattern metrics (saturation, brightness, hue, and percent gray patch) for red-eyed treefrogs (*Agalychnis callidryas*) sampled from 12 sites along the Pacific coast of Costa Rica. Individuals are color coded by region (see Supplemental Figure 6 for legend).

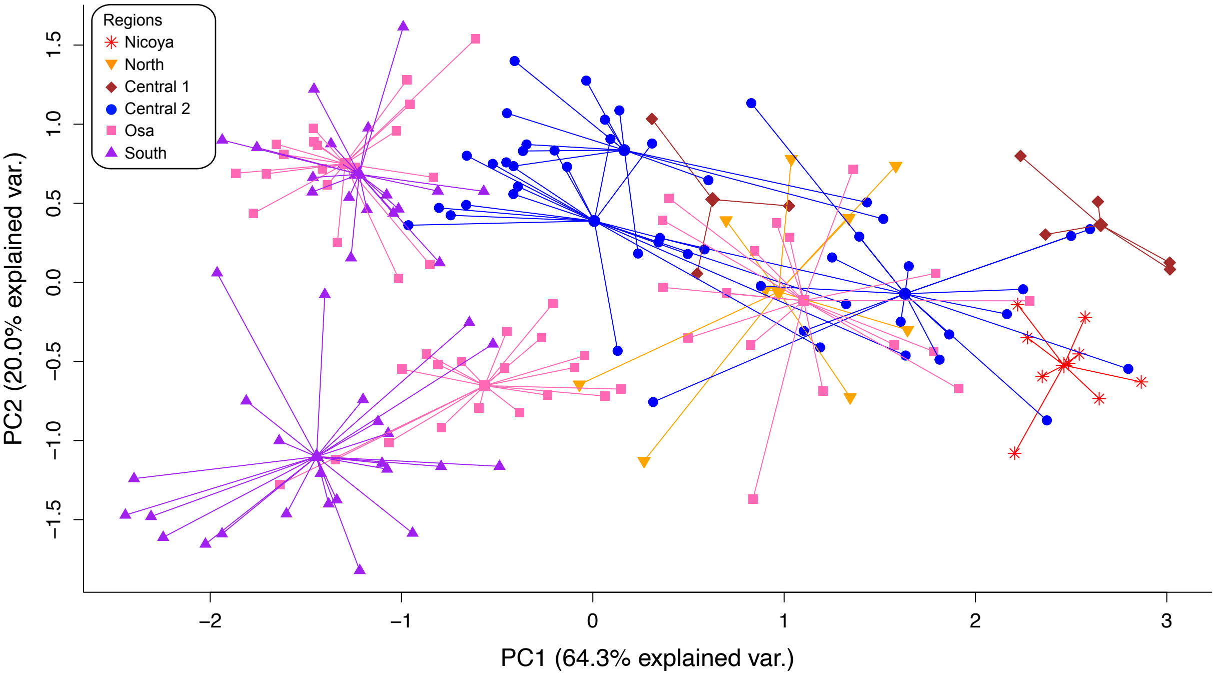


Supplemental Figure 6: Principal component analysis of leg coloration measurements (saturation, brightness, and hue) for red-eyed treefrogs (*Agalychnis callidryas*) sampled from 12 sites along the Pacific coast of Costa Rica. Lines connect individuals (small dots) with the mean phenotype of each sampling site (large dots). Individuals and means are color coded by region.

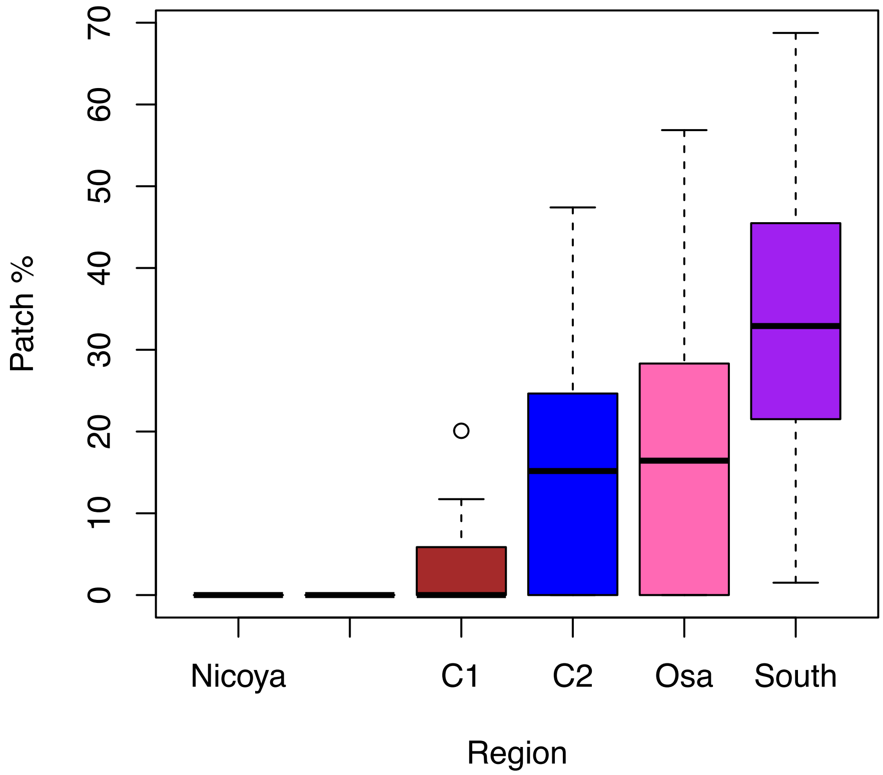


Supplemental Figure 7: Boxplot showing the distribution of the percent of hind leg taken up by the inner blue/gray patch for red-eyed treefrogs (*Agalychnis callidryas*) sampled from 12 sites along the Pacific coast of Costa Rica. Horizontal bar shows the average percent patch for each region.

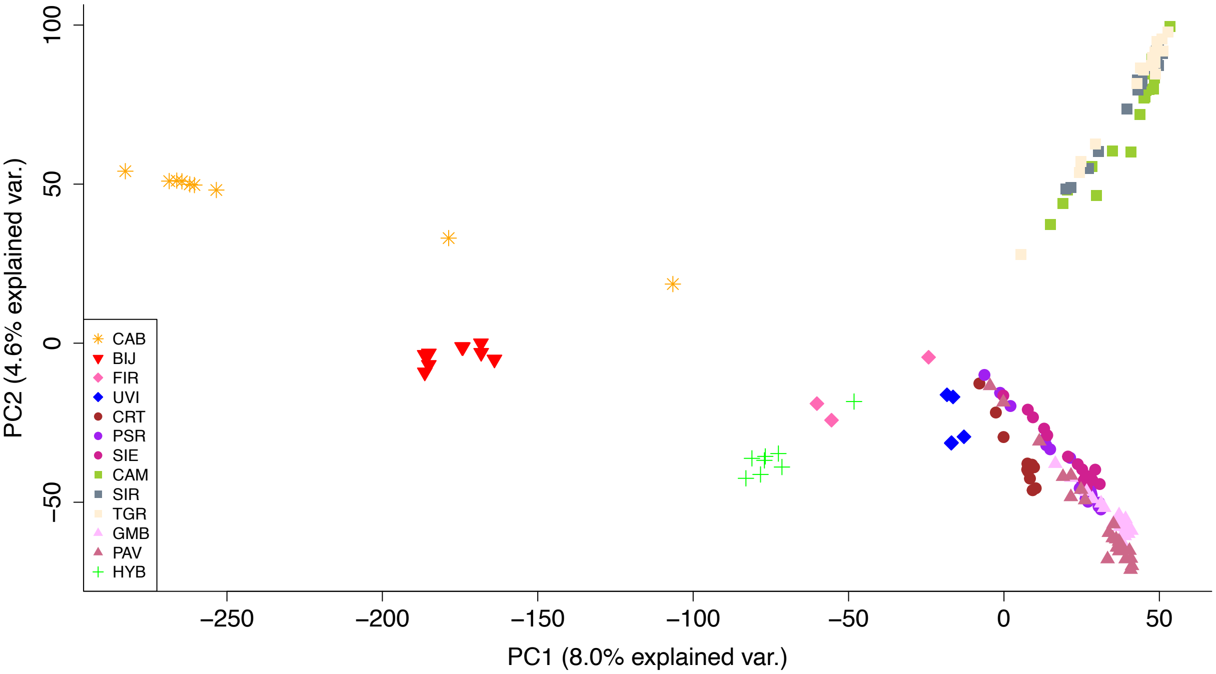


Supplemental Figure 8: Principal component analysis of genomic SNPs for red-eyed treefrogs (*Agalychnis callidryas*) sampled from 12 sites along the Pacific coast of Costa Rica. Symbols represent individuals. Symbol type represents region, while color represent sampling site. Green plus signs are lab generated hybrids between Site 2 and Site 12.

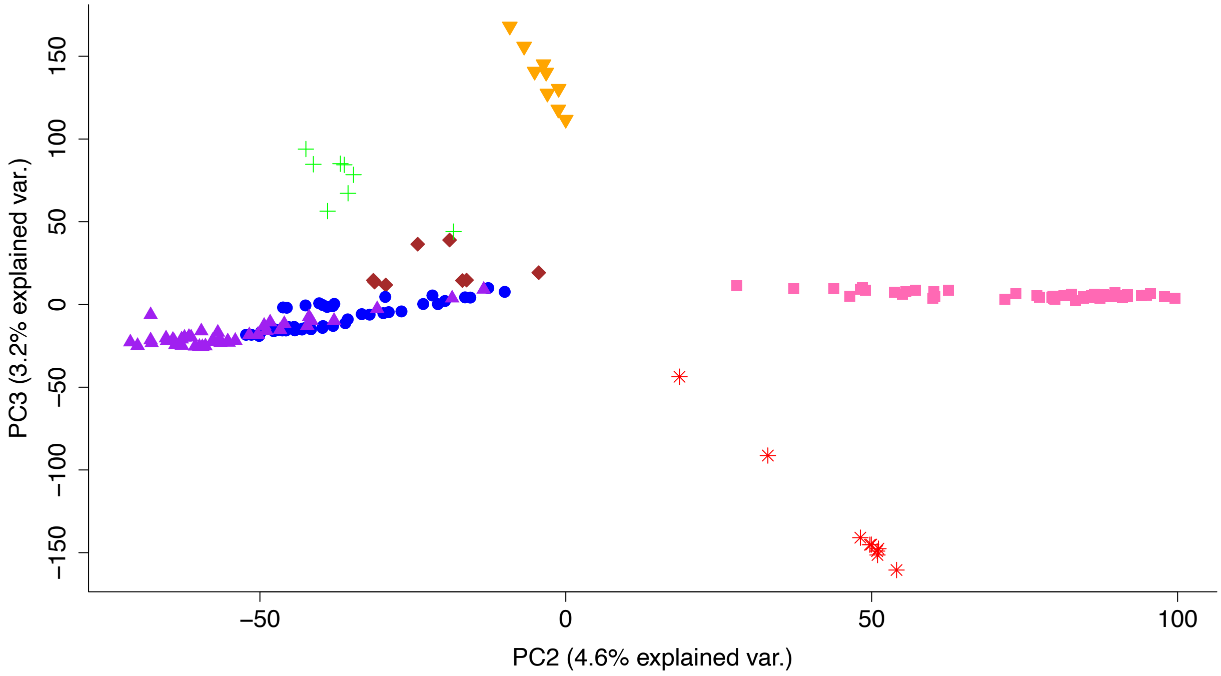


Supplemental Figure 9: Principal components 2 and 3 from principal components analysis of genomic SNPs for red-eyed treefrogs (*Agalychnis callidryas*) sampled from 12 sites along the Pacific coast of Costa Rica. Symbols represent individuals. Both type of symbol and color represent region (see Supplemental Figure 6 for legend). Green plus symbols are lab generated hybrids between Site 2 and Site 12.

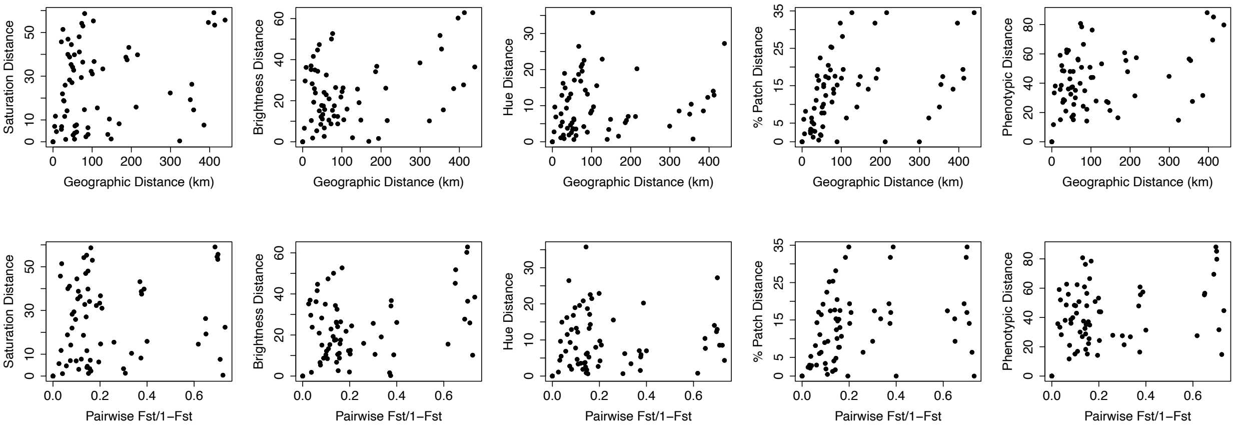


Supplemental Figure 10: Relationships between over-land phenotypic measurements and overland geographic or genetic distance for 12 sites of red-eyed treefrogs (*Agalychnis callidryas*) along the Pacific coast of Costa Rica. Mantel tests found no significant relationships between any comparisons.

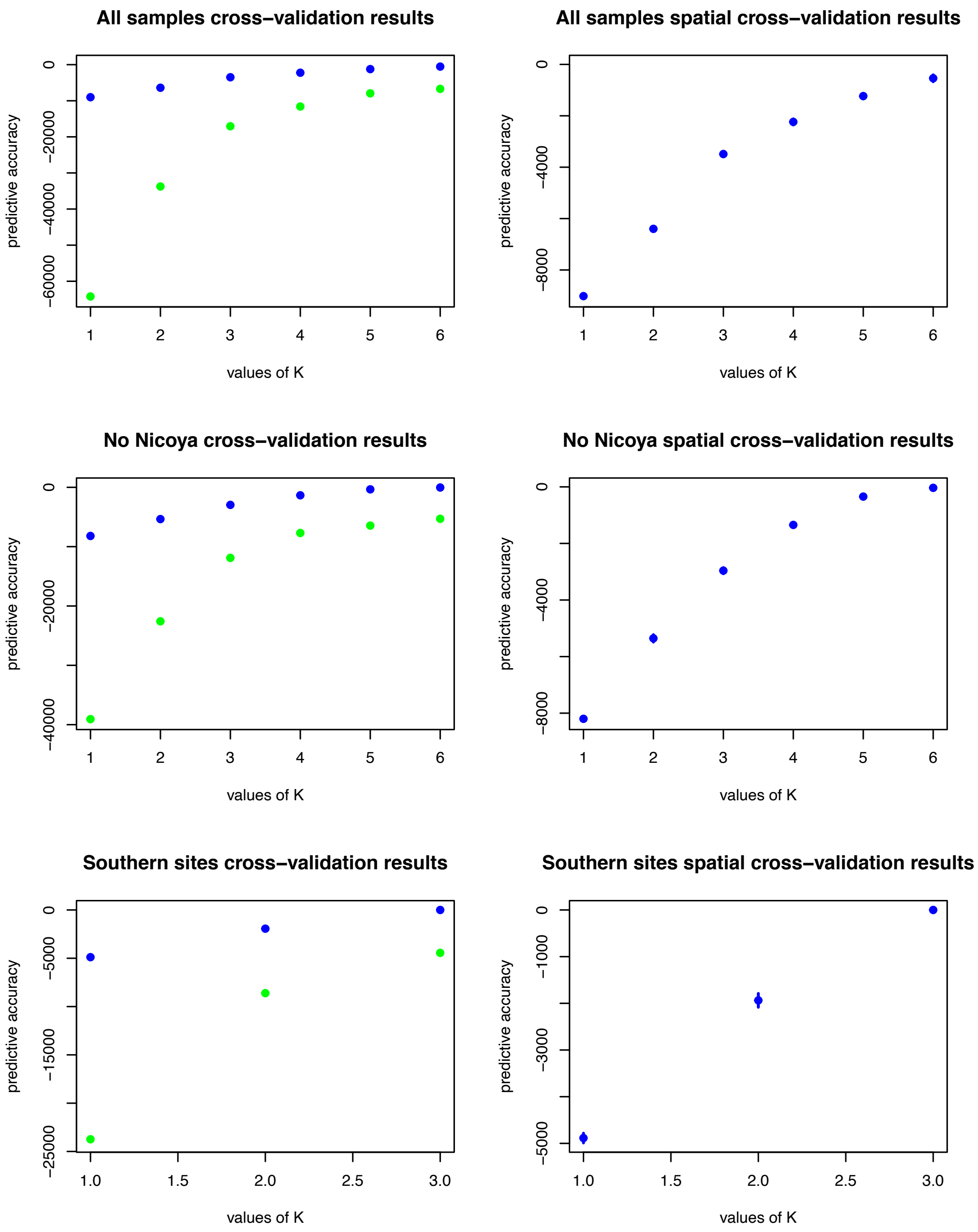


Supplemental Figure 11. Scatterplot comparison of the predictive accuracy of *K*-values from conStruct cross-validation analysis of three different data subsets, each run with eight replicates. Green and blue dots represent nonspatial and spatial models, respectively. All spatial models had higher predictive accuracy than nonspatial models.

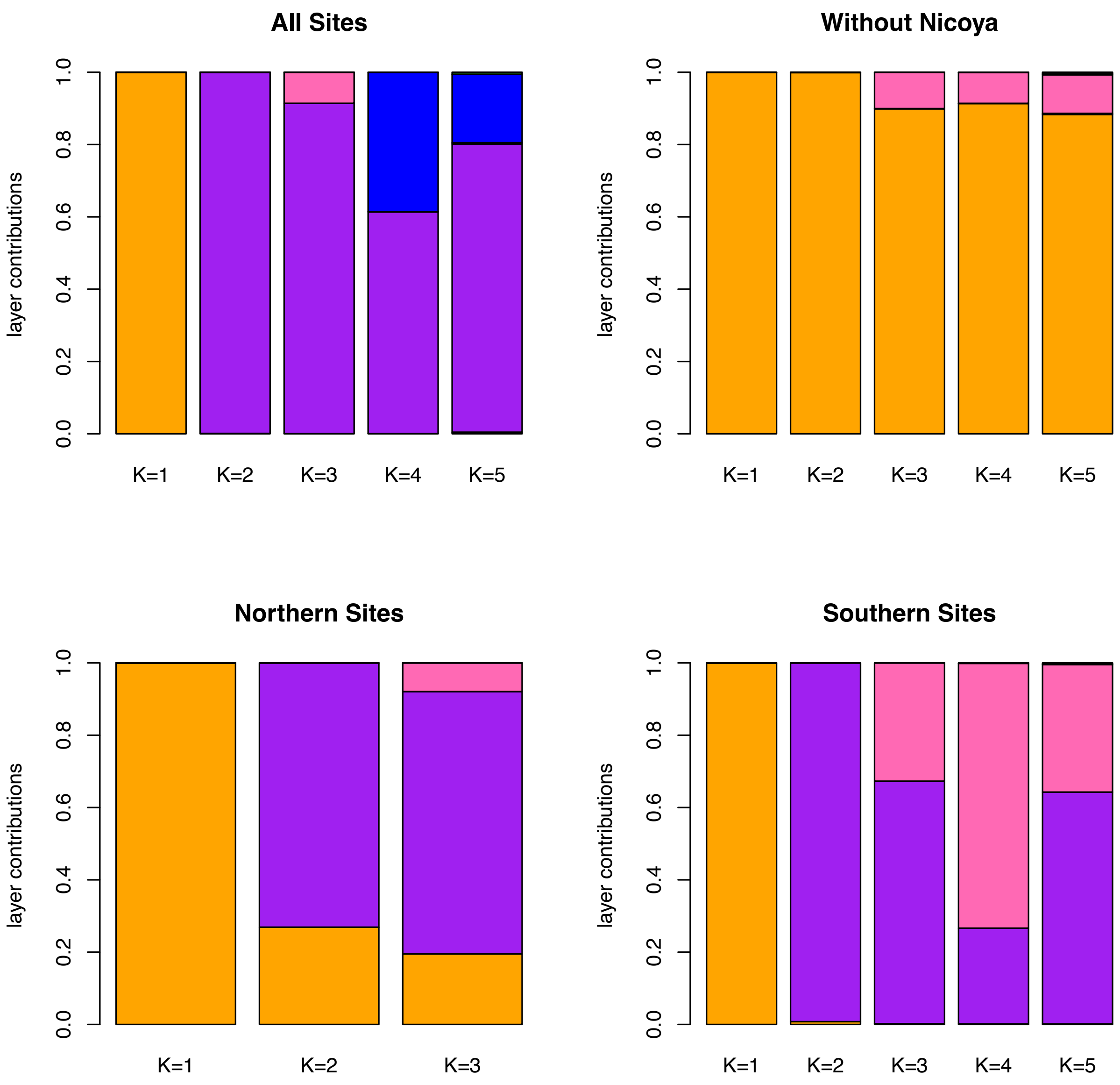


Supplemental Figure 12. Barplot of the contribution of each layer to parametric covariance for K-values 1 through 5 from spatial conStruct analyses. Colors show proportions of different layers.

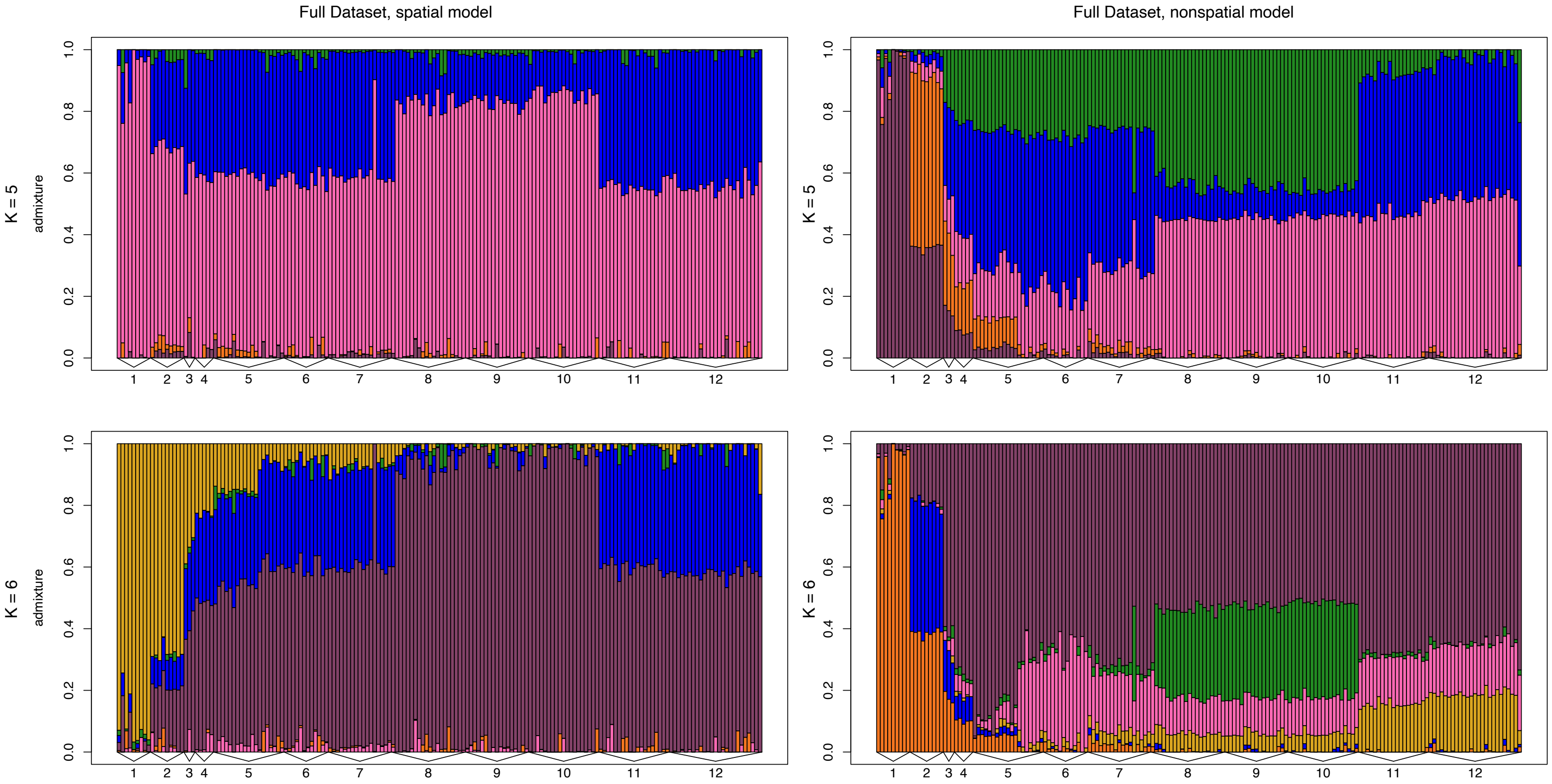


Supplemental Figure 13. Barplots of admixture proportions for red-eyed treefrogs (*Agalychnis callidryas*) sampled from 12 sites along the Pacific coast of Costa Rica. Admixture proportions shown were generated using conStruct for *K* = 5–6 for both non-spatial and spatial models. Each bar represents an individual. The color of the bar shows the proportion of the individual's genome that is assigned to each of *K* layers.

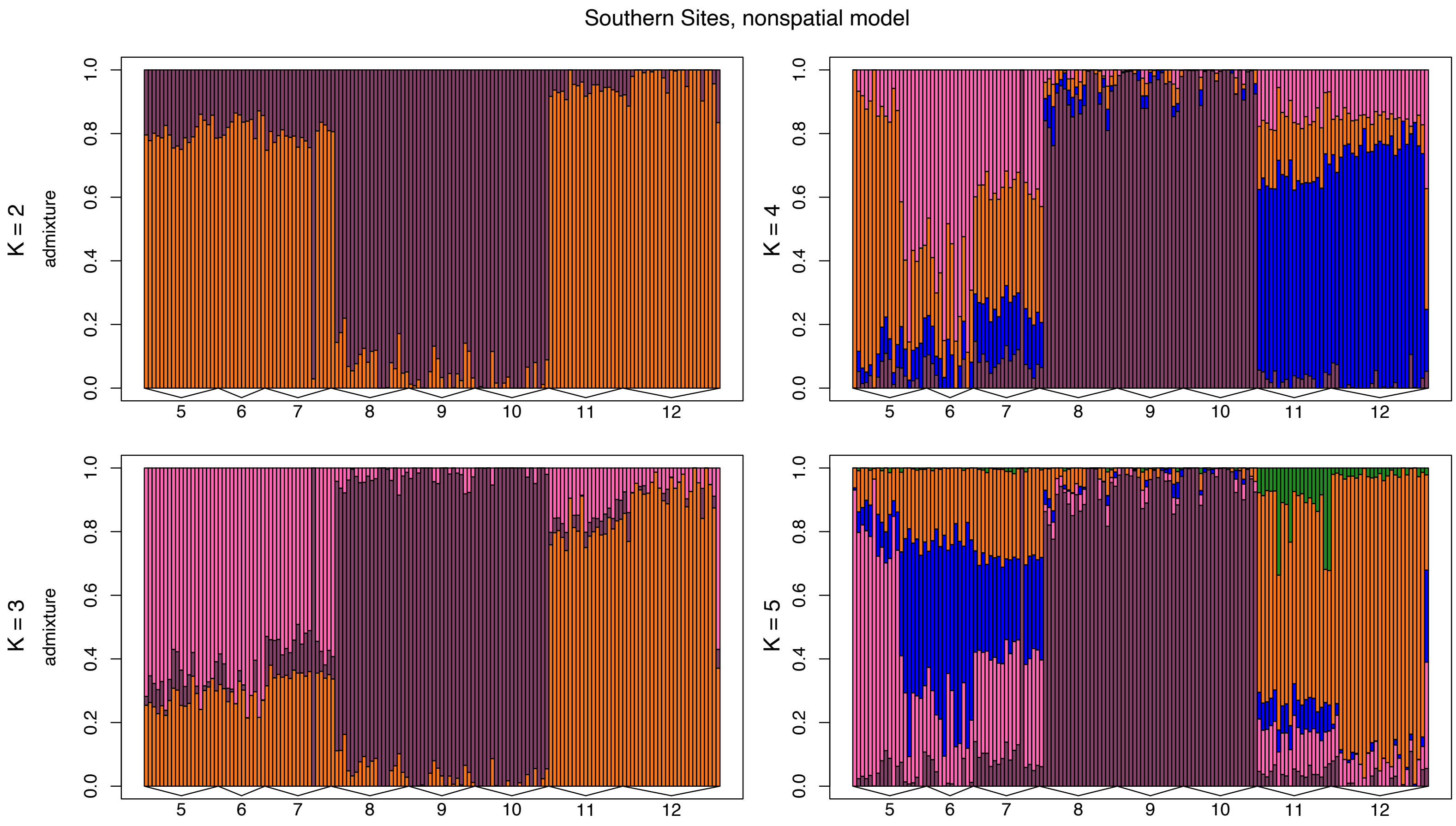


Supplemental Figure 14. Barplots of admixture proportions for red-eyed treefrogs (*Agalychnis callidryas*) sampled from southern sites (Sites 5-12). Admixture proportions shown were generated using conStruct for $K = 2–5$ for nonspatial models. Each bar represents an individual. The color of the bar shows the proportion of the individual's genome that is assigned to each of $K$ layers.

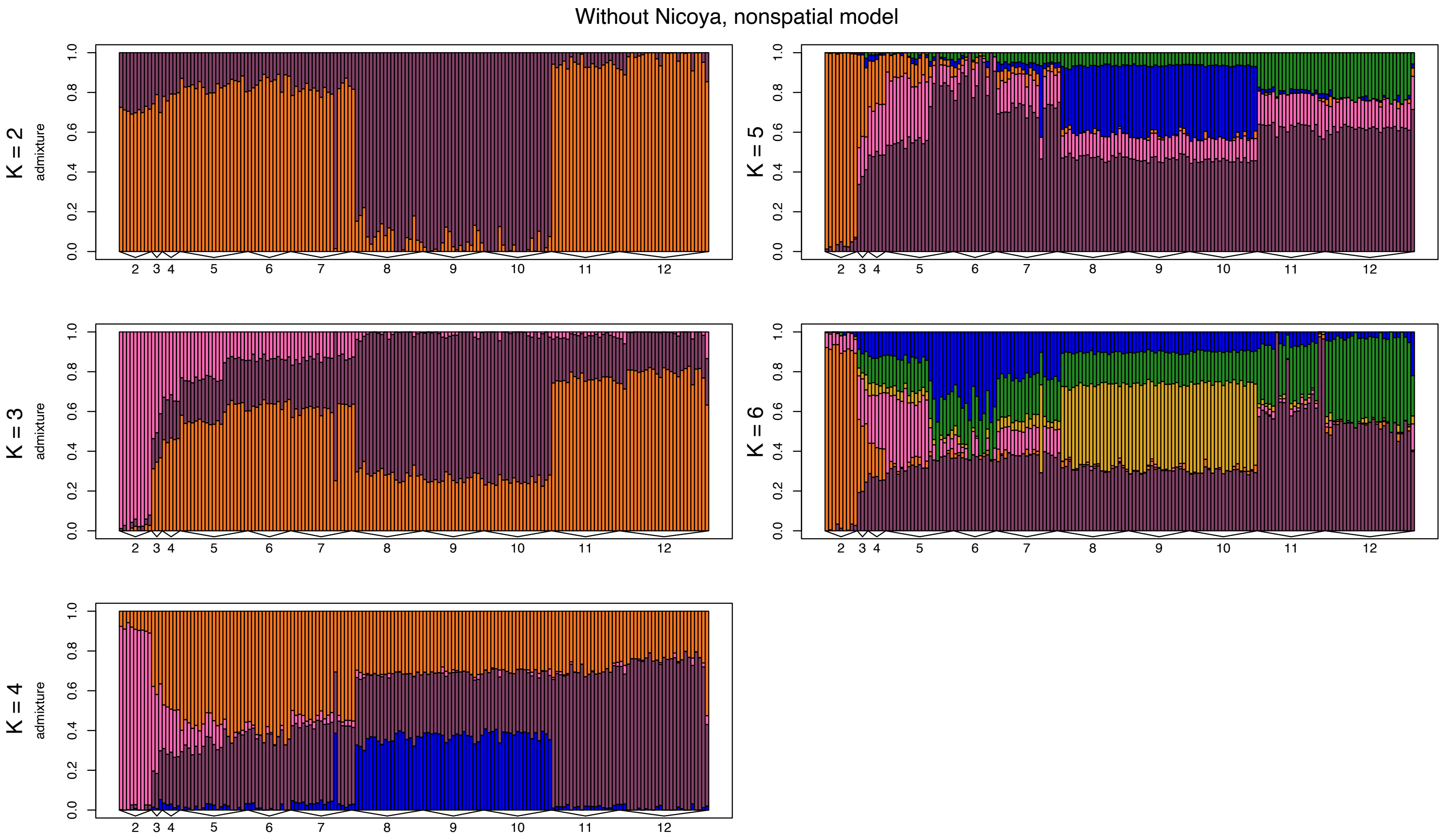


Supplemental Figure 15. Barplots of admixture proportions for red-eyed treefrogs (*Agalychnis callidryas*) sampled from a subset of the dataset that includes Sites 2–12. Admixture proportions shown were generated using conStruct for *K* = 2–4 for spatial models. Each bar represents an individual. The color of the bar shows the proportion of the individual's genome that is assigned to each of *K* layers.

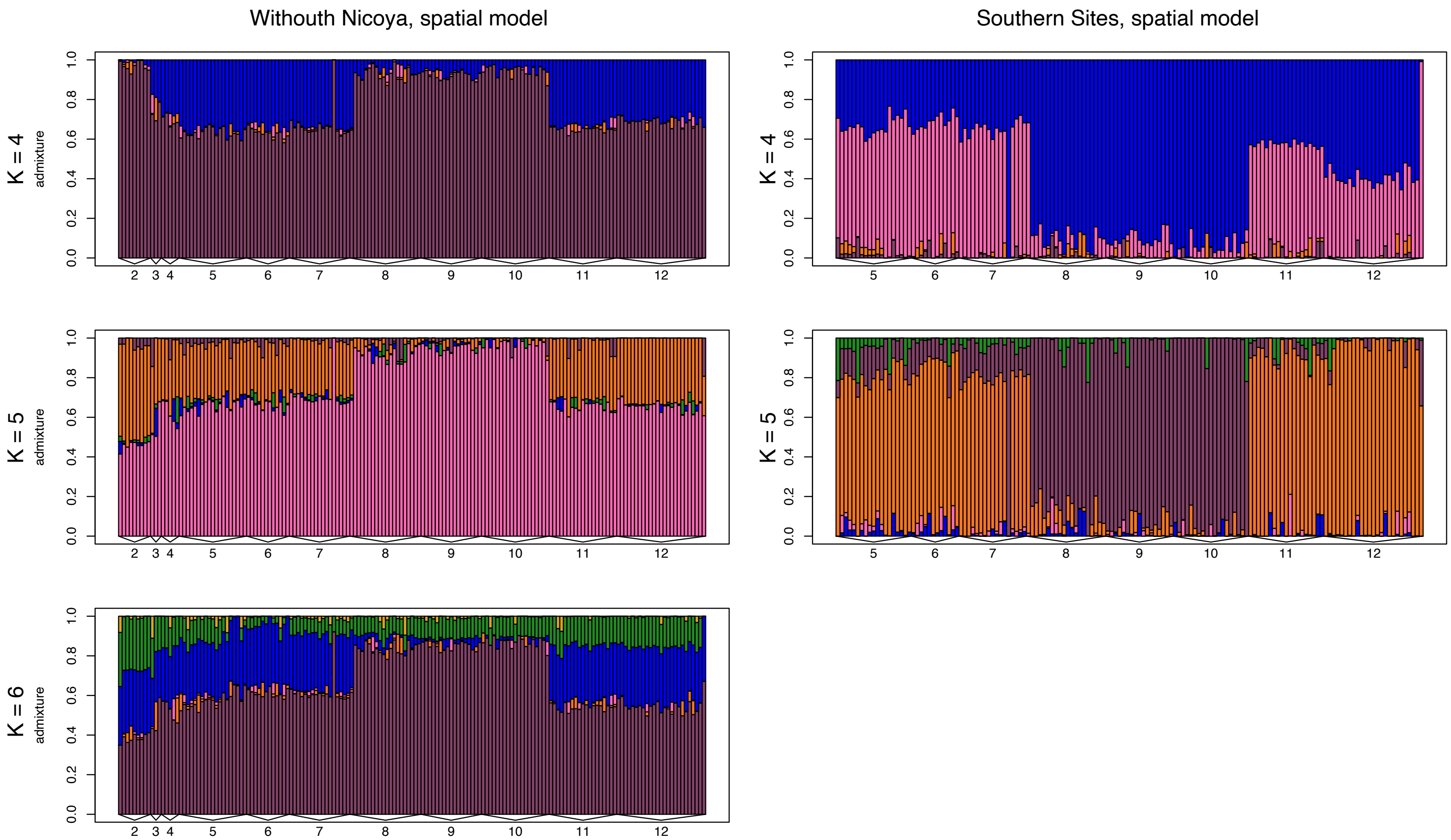


Supplemental Figure 16. Barplots of admixture proportions for red-eyed treefrogs (*Agalychnis callidryas*) sampled from two data subsets: without Nicoya (Sites 2–12) and southern sites (5-12). Admixture proportions shown were generated using conStruct for $K$ = 4–6 or K = 5–6 respectively for spatial models. Each bar represents an individual. The color of the bar shows the proportion of the individual's genome that is assigned to each of $K$ layers.

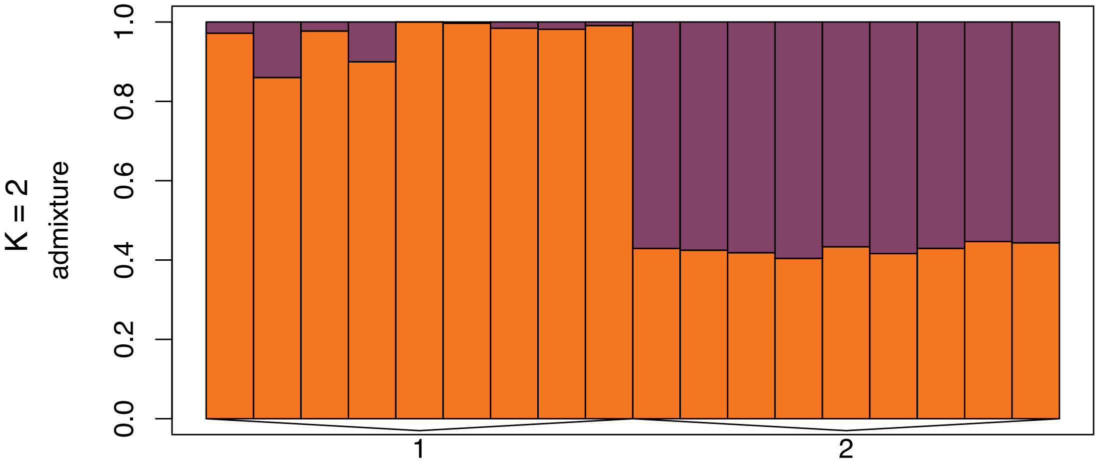


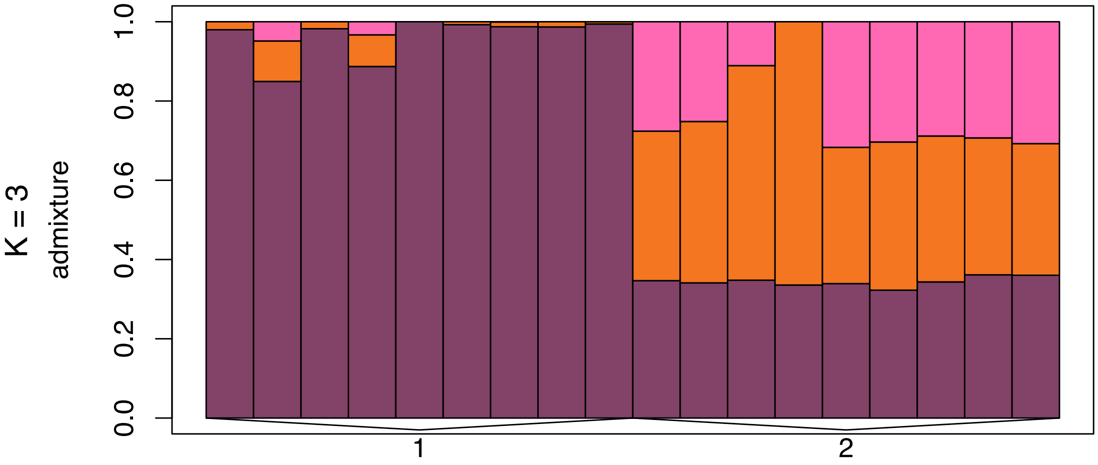


Supplemental Figure 17. Barplots of admixture proportions for red-eyed treefrogs (*Agalychnis callidryas*) sampled from northern sites (Sites 1–2). Admixture proportions shown were generated using conStruct for $K$ = 2–4 for nonspatial models. Each bar represents an individual. The color of the bar shows the proportion of the individual's genome that is assigned to each of $K$ layers.

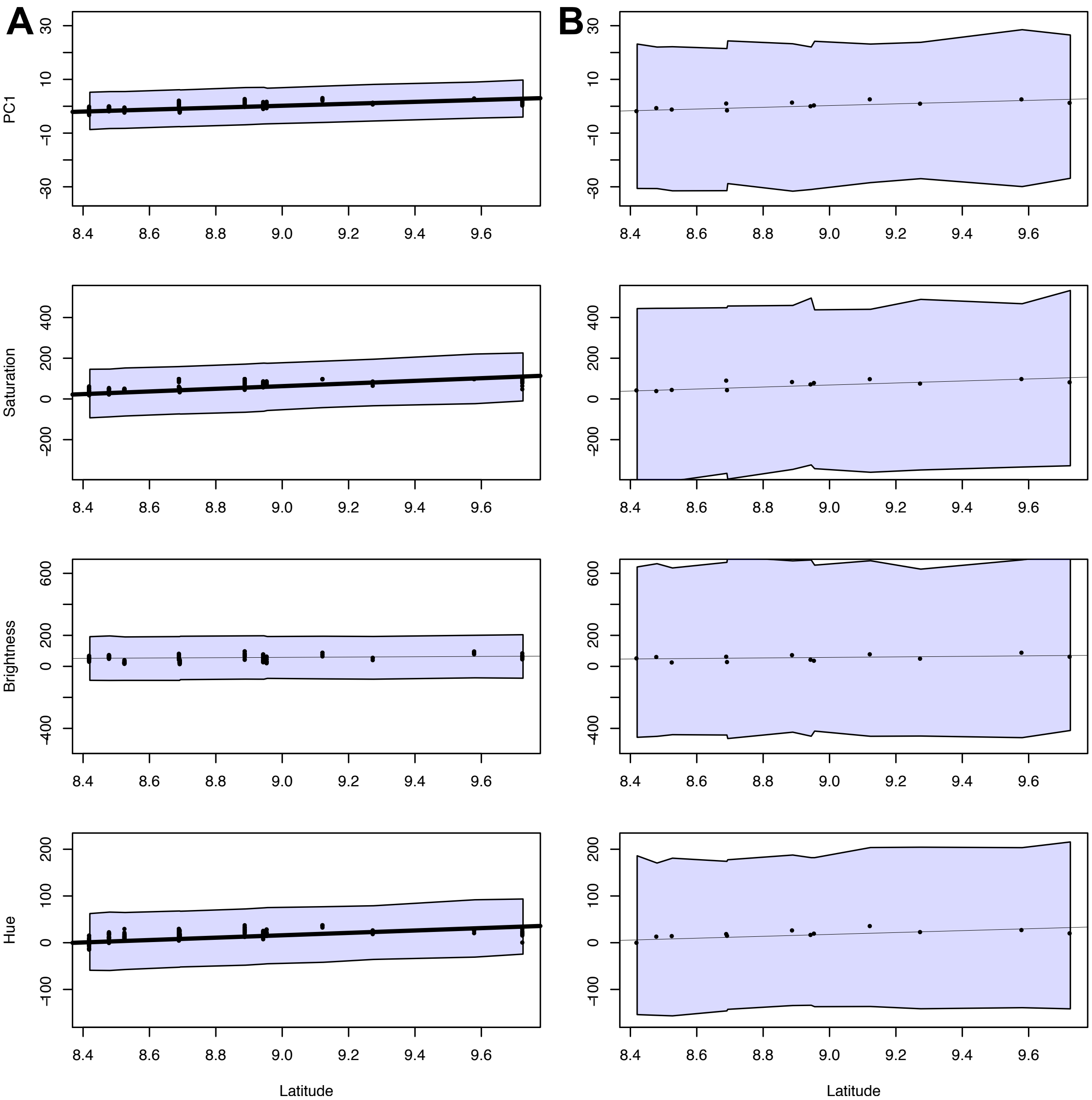


Supplemental Figure 18: Results of Bayesian models with Dunn-Bonferroni corrected credible intervals (CI) showing the relationships between phenotype measurements (PC1, saturation, brightness, and hue) and latitude for (A) individuals and (B) site averages. Bolded line indicates that the 95% CI for the parameter estimate did not overlap with zero.

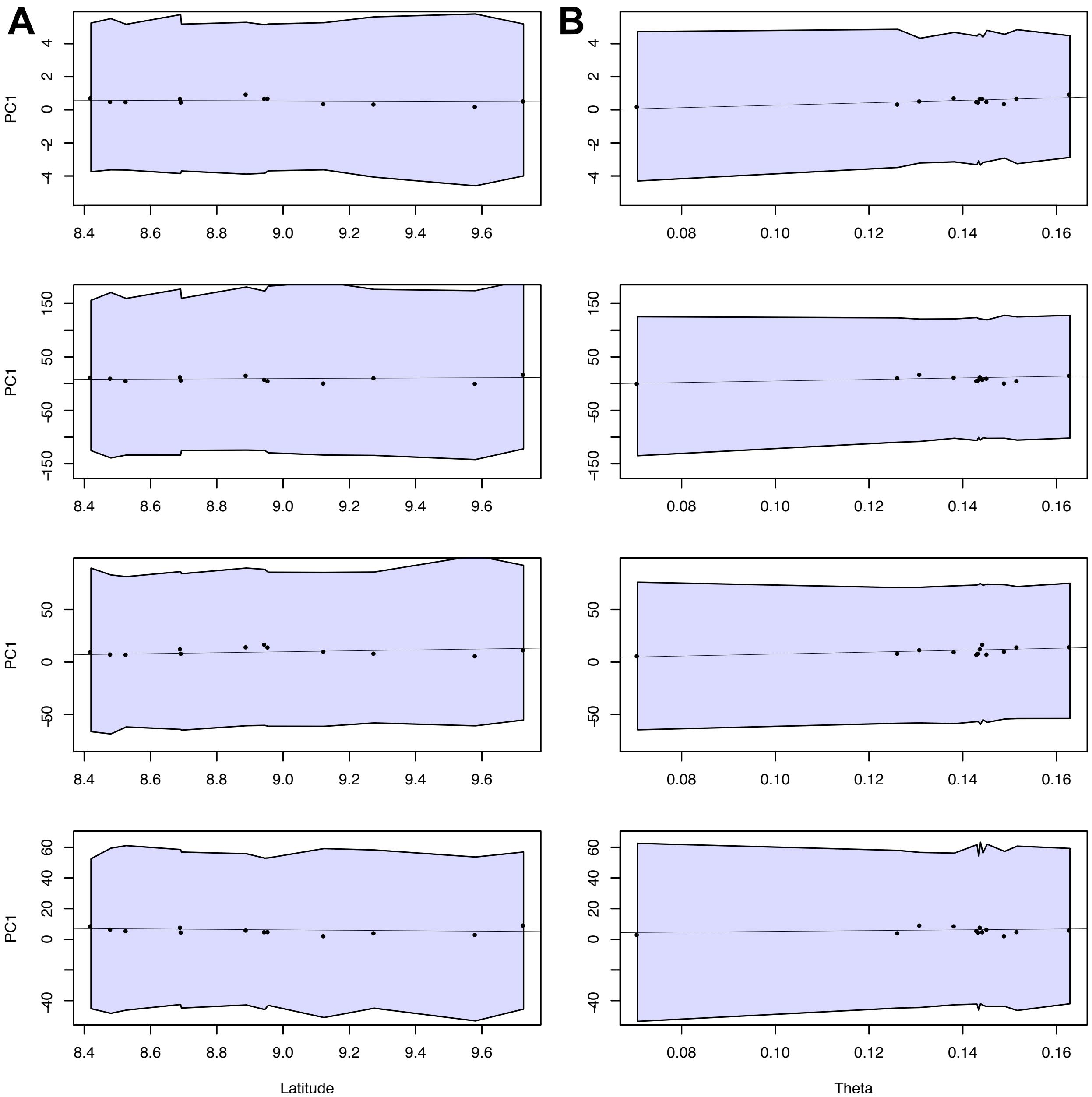


Supplemental Figure 19: Results of Bayesian models with Dunn-Bonferroni corrected credible intervals (CI) showing the relationships between (A) variation in phenotype measurements (PC1, saturation, brightness, and hue) and latitude for individuals and site averages, and (B) variation in phenotype measurements and Wu and Watterson's theta.
